# Towards Generalizable Protein-ligand Co-folding with ACER

**DOI:** 10.64898/2026.06.02.728568

**Authors:** Nopsinth Vithayapalert, Francesca Grisoni

## Abstract

Predicting protein–ligand complex structures is a central challenge in drug discovery. While recent co-folding models such as AlphaFold-3 achieve accurate structure prediction, they fail to generalize to underexplored binding interfaces – systematically misplacing ligands, particularly for allosteric or structurally novel targets. To address this gap, we present **ACER** (**A**daptive **C**o-folding via pocket **E**xploration and pose **R**anking), a training-free framework that (a) enables co-folding models to systematically explore alternative binding pockets, and (b) leverages the discovered pockets to increase pose accuracy. Our method enables the efficient discovery of non-prevalent pockets without prior expert knowledge. ACER improves pocket discovery and pose accuracy on allosteric targets and structurally novel complexes, successfully modeling binding interfaces that are under-represented or absent from the training set. Our results demonstrate how improved sampling dynamics enhance the generalisability of co-folding models without retraining.

## 1 Introduction

Characterizing protein-ligand interactions at the structural level provides a fundamental understanding of how a small molecule modulates the biological function of one or more macromolecular targets (Zhao et al., 2022; Ciulli, 2013). Traditional structural biology techniques, *e.g*. X-ray crystallography and cryo-electron microscopy, can accurately resolve the structure of protein-ligand complexes, but they remain time-consuming, costly, and difficult to scale for high-throughput applications (Shi, 2014). Co-folding models have recently emerged as scalable computational tools to predict the structures of protein-ligand complexes and complement traditional experimental techniques (Abramson et al., 2024; team et al., 2024; OpenFold3 Team, 2024; Zhang et al., 2026; Passaro et al., 2025). Their competitive performance in both structure prediction and molecular docking (Durairaj et al., 2024) highlights their potential to enable high-throughput virtual screening and guide molecule design.

Despite these advances, co-folding models exhibit a critical failure mode (Škrinjar et al., 2025b): their tendency to strongly memorize the training data. The model’s predictive power highly correlates with the similarity between training and test samples, as further evidenced by the fact that models misplace allosteric ligands into orthosteric binding sites for protein targets whose allosteric sites are underrepresented (Nittinger et al., 2025; Parikh et al., 2026). Similarly, protein conformations are strongly affected by the ratio of holo- and apo-structures in the training set (Yu et al., 2026). These findings suggest that co-folding models have learned the template (’memorization’) rather than the underlying physics of protein-ligand interactions.

One of the main bottlenecks of existing co-folding models is the incorrect ligand placement in the protein pocket, which is estimated to occur in allosteric complexes (Parikh et al., 2026; Nittinger et al., 2025) and in proteins that are underrepresented in the training data (Yu et al., 2026; Škrinjar et al., 2025b). Correct ligand placement is relevant, as most of the targets in drug discovery campaigns are structurally novel. This is particularly consequential for allosteric targets, where the functional pocket is often transient and only forms upon ligand-induced conformational change. This pocket memorization behavior therefore represents a critical gap in co-folding models.

To overcome this bottleneck, explicit pocket constraints have been proposed as guidance for cofolding (Passaro et al., 2025; Zhang et al., 2026). However, this technique requires prior knowledge about the correct pocket, which is usually unavailable for novel or out-of-distribution targets. Existing pocket identification tools (Le Guilloux et al., 2009; Krivák & Hoksza, 2018) predict binding-site residue probabilities in the absence of a ligand, which might miss ligand-induced conformational rearrangements Meller et al. (2023). In this context, improving pocket discovery for novel targets holds untapped potential on the path to generalizable co-folding models.

In this work, we present **ACER** (**A**daptive **C**o-folding for pocket **E**xploration and pose **R**anking), a framework aimed at improving the capacity of co-folding models to discover and prioritize binding pockets – ultimately to retrieve more accurate protein-ligand structures. Our main contributions are:

1. **Inference-time pocket exploration**, which expands the search for binding pockets without retraining or major architectural modification. We implement two complementary strategies to enable the exploration the possible pocket landscape: (1) decoy ligand conditioning, which augments the input representation with auxiliary ligands that occlude default binding pockets; (2) iterative pocket repulsion, which applies negative gradients via Feynmann-Kac (FK) steering (Singhal et al., 2025). These strategies were implemented and tested on Boltz-2 (Passaro et al., 2025) and Protenix-v1 (Team et al., 2026).
2. **Ensemble-based ranking**, aimed at improving the prioritization of predicted ligand poses. This is achieved via normalized scores that derive from protein-ligand conformational ensembles, predicted under each pocket constraint.
3. **Improved generalization on hard co-folding benchmarks**. ACER successfully places ligands into experimentally observed binding pockets for allosteric compounds in those cases when state-of-the-art co-folding mistakenly places ligands into orthosteric pockets. When tested on the challenging subset of the Runs N’ Poses benchmark (Škrinjar et al., 2025a) – where binding interfaces are highly dissimilar to those in training set – this approach substantially outperforms the co-folding baselines.

## 2 Background and Related Works

### Generative models for biomolecular interactions

Co-folding is built on conditional diffusion governed by two core components (Passaro et al., 2025; Abramson et al., 2024): (1) *trunk representations* that encode geometric features, and (2) a *denoiser* that iteratively recovers the clean structure from noise. Concretely, the model tokenizes protein sequences at the residue level and ligands at the atom level, yielding *N* = *N*_P_ + *N*_ℓ_ tokens (protein residues and ligand atom tokens, respectively). The trunk produces a single representation **s** ∈ ℝ^*N* ×*d*^ and a pair representation **z** ∈ ℝ^*N* ×*N* ×*d*^, which together form the conditioning variable **Z** = (**s, z**) for the denoiser. Recent models (Passaro et al., 2025) augment the sampling procedure with training-free Feynmann-Kac (FK) steering via Sequential Monte-Carlo (SMC), applying gradient updates to impose physico-chemical constraints. Yet, FK steering has been used to drive sampling toward a known pocket, which renders it unsuitable for novel targets or allosteric complexes. While recent works applied Twisted Diffusion (Richman et al., 2026) and latent perturbation (Lee et al., 2026) to explore protein conformational landscapes, its potential for *exploring* alternative distinct ligand binding pockets remains untapped.

### Memorization in co-folding

Protein-ligand co-folding models (Abramson et al., 2024; Passaro et al., 2025; OpenFold3 Team, 2024; Zhang et al., 2026) achieve state-of-the-art performance on structure-prediction and blind-docking benchmarks (Škrinjar et al., 2025b; Durairaj et al., 2024). Yet, accuracy drops as the targets diverge from ‘familiar’ biochemical space, and co-folded structures are sensitive to mutations at key binding-site residues (Masters et al., 2025). These findings point to strong memorization of the training data rather than genuine learning of biomolecular interaction physics – a fundamental issue for which the field lacks a principled solution.

### Pocket identification

Conventional methods (Le Guilloux et al., 2009; Krivák & Hoksza, 2018) detect the pockets directly from protein structures in the absence of ligands, while more recent tools (Wang & Dokholyan, 2025) incorporate chemical probes and learned molecular fingerprints. All share the limitations of co-folding models: their accuracy drops over distribution shifts, e.g., for allosteric sites or unseen pockets where co-folding fails (Parikh et al., 2026). Naively combining the two, therefore, inherits rather than solves the generalization problem. Advances in pocket identification have been reported (Isomorphic Labs Team, 2026); however, the closed-source nature precludes any deeper analysis of gains in co-folding generalizability.

### Pose ranking in molecular docking

Search-based methods such as AutoDock Vina (Trott & Olson, 2010) rely on physics-based scoring functions, treating pocket side-chains as rigid and ignoring ligand-induced fit effects. Co-folding instead ranks structures via confidence scores, which do not necessarily correlate with the success rate (Team et al., 2025). Recently, Molecular Dynamics (MD) has been combined with Markov state models to surface cryptic and allosteric pockets that are only visible in multi-conformational states (Biswas et al., 2025), but at the cost of expensive simulations.

## 3 Method

ACER addresses ligand misplacement in co-folding models through a two-stage framework (Fig. 1a): *(1) binding pocket exploration* and (2) *pose ranking* via an ensemble approach.

**Figure 1:**
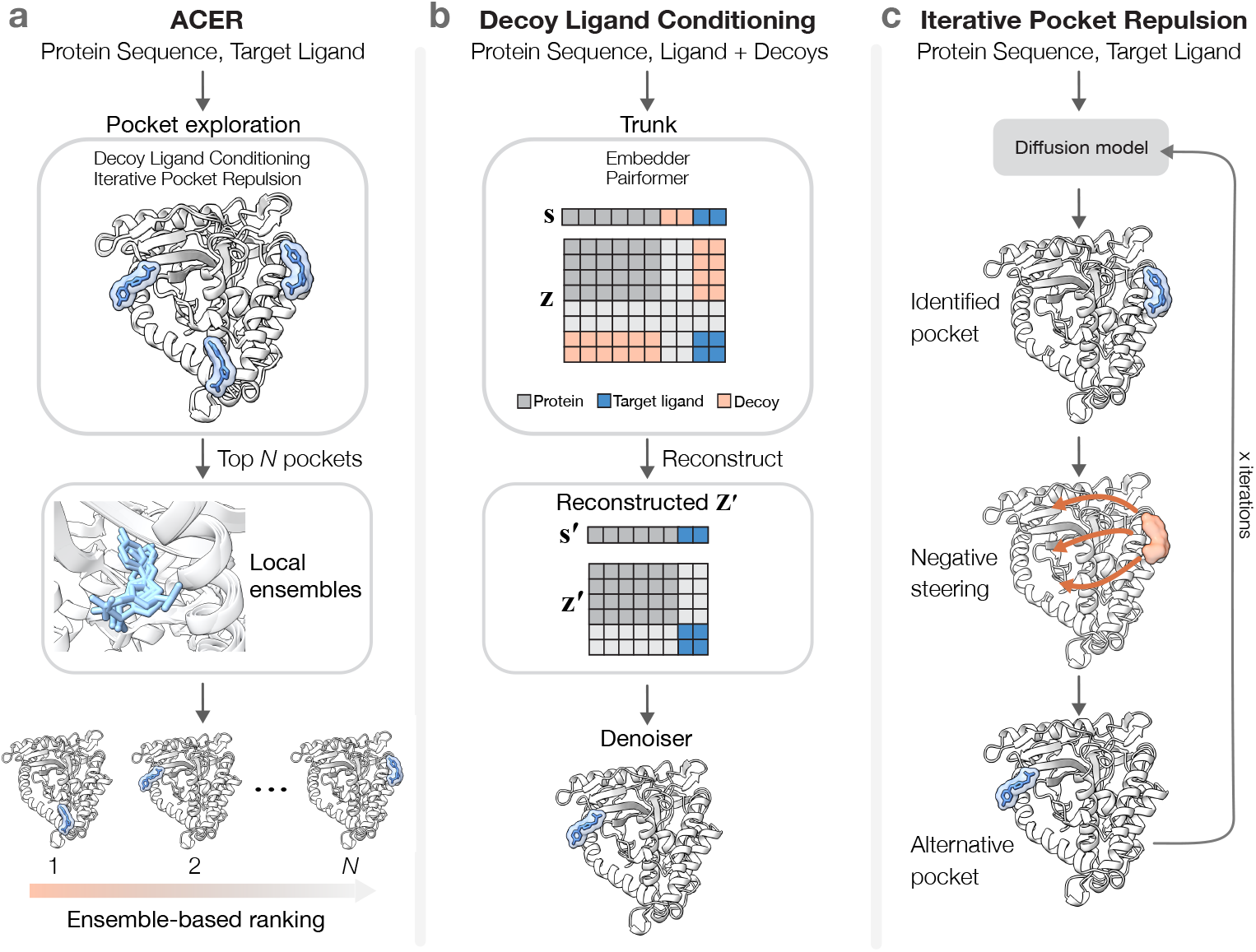
Overview of ACER. (Adaptive Co-folding for pocket Exploration and pose Ranking). **(a)** ACER co-folds complexes across several candidate binding pockets. Each pocket then serves as a constraint for generating a local ensemble, from which top-ranked poses are selected using ensemble-based scores. **(b)** Given a protein *P* and target ligand ℓ, one or more decoy ligands 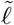 are introduced as additional trunk inputs. After processing by the Pairformer, the decoy tokens are discarded to reconstruct conditioning variable that co-folds *P* and ℓ into alternative pockets. **(c)** Inference-time guidance to direct the ligand placement away from the previously explored pockets. At each round, the previously explored pocket is added to a repulsion set *ℬ*, steering the denoiser away from previously explored sites.

### 3.1 Binding pocket exploration

We propose two strategies: (1) *decoy ligand conditioning* (Fig. 1b), which augments trunk inputs with auxiliary ligands to redirect the model’s distributional bias away from dominant pockets, and *(2) pocket repulsion* (Fig. 1c), which steers the denoising trajectory away from previously sampled binding sites. The former leverages the model’s implicit learning of interactions to explore alternative pockets in a parameter-free manner; the latter provides specific, user-defined steering. Together, they expand the effective search space of binding sites without retraining or architectural modifications. By its design, ACER requires a co-folding model that supports inference-time steering such as Boltz-2 (Passaro et al., 2025), Protenix-v2 (Zhang et al., 2026), or OpenBind-v0 ().

### 3.1 Decoy ligand conditioning

Motivated by the evidence that co-folding with two identical allosteric ligands increases the likelihood of placing one into the correct pocket (Nittinger et al., 2025), we introduce *decoy ligand conditioning* (Fig. 1**b**). Given a protein *P* and a ligand ℓ, we augment the trunk inputs with one or more decoy ligands 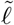 (comprising 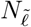 atom tokens). These decoys can be molecules identical to ℓ or generic chemical probes. The augmented inputs are processed by the Pairformer module to compute cross-token interactions between all inputs, producing the conditioning representations 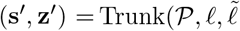, where 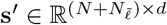 contains token-level representations enriched by cross-token interactions, and 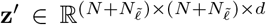 encodes pairwise interactions across all token pairs, including those involving 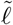. We then reconstruct the conditioning variable by discarding all rows and columns corresponding to 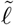 tokens, yielding:

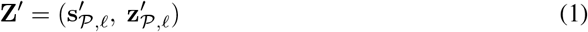

where 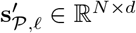 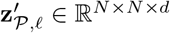 have the same dimensions as **Z** but are enriched by decoy-induced cross-token interactions. The denoiser generates structures conditioned on this reconstructed variable, **x**_0_ ~ *p*_*θ*_(**x** | **Z**′), such that 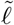 influences only the conditioning representations but does not participate in generation during denoising. Intuitively, 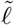 occludes the dominant pocket in **z**′, reducing its dominant signals in **Z**′, and freeing ℓ to explore alternative binding sites. We verify this occlusion mechanism empirically in Appendix B.1, where decoy ligand conditioning shifts both the trunk’s predicted contacts and the denoiser’s attention away from the orthosteric pocket and toward alternative sites.

We use four decoys that are identical to the considered ligand – native ligands. To assess whether non-native binders could improve pocket discovery, we additionally ablated the choice of decoy ligand type using three ‘frequent hitters’ chemical probes (Fig. D5), binding molecules that bind to diverse proteins across the Protein Data Bank Wang & Dokholyan (2025). Non-native binders offered no clear advantage over native ligands for pocket exploration.

#### 3.1.2 Iterative pocket repulsion

While decoy ligand conditioning influences representation learning, inference-time steering is a plugin strategy for controlling generation to respect specific constraints. To complement the FK-steering framework of Boltz-2 – which accounts for physico-chemical constraints (Passaro et al., 2025) – we introduce a *pocket repulsion potential*, which biases the denoising trajectory away from a specified pocket (via negative gradients), toward alternative binding contacts.

We penalise each ligand atom for being closer than a minimum distance (*d*_min_) to the surface of a virtual sphere of radius *r*_*g*_ centred on the pocket anchor **c**_*g*_, where *g* denotes the pockets to steer away from. The pocket anchor **c**_*g*_ is computed as the mean centroid of its constituent residue tokens *t*:

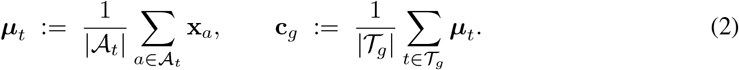

where *A*_*t*_ is the set of atoms in *t*, **x**_*a*_ ∈ ℝ ^3^ is the position of atom *a*, and *T_g_* denotes residues within 4 Å from the ligand atoms comprising pocket *g*.

The repulsion potential is then defined over all *G* pockets and all ligand atoms *A*_lig_, penalising any ligand atom that lies within a distance *d*_min_ of the surface of the sphere centred on **c**_*g*_ with the hyperparameter of virtual pocket radius *r*_*g*_:

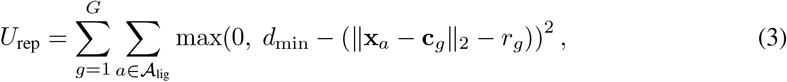

We apply these potentials iteratively. Each inference round collects the occupied pocket as a new anchor, which is then added to the repulsion set *B* for all subsequent rounds (Algorithm 1; full parameter settings in Appendix A).

### 3.2 Ensemble-based ranking

Once putative pockets have been identified, it is crucial to distinguish correctly docked poses from geometrically plausible but non-native ones (*i.e*., those not reflecting the experimentally observed binding mode). Co-folding models generate a single static pose per run, which may miss the correct binding mode due to the intrinsic flexibility of protein-ligand complexes and the dynamics of ligand binding. We therefore generate local conformation ensembles around each candidate pocket — leveraging the fact that Boltz-2 was trained on MD simulations — and rank poses by their consistency with the local ensemble. For each pocket *p*, we collect *n* pose scores {*s*_1_, ‖, *s*_*n*_} to compute the z-score i.e. *z*_*i*_ = (*s*_*i*_ − *µ*_*p*_)*/σ*_*p*_ where *i* = 1, ‖, *n* indexes the poses within pocket *p, µ*_*p*_ and *σ*_*p*_ are the mean and standard deviation of pose scores within *p*.

We then weight each score by its proximity to the mean, either via a Gaussian or Lorentzian normalization, both of which peak at the ensemble mean and decay for outliers. The *Gaussian* weight penalises deviations sharply, whereas the *Lorentzian* weight down-weights outliers more gradually:

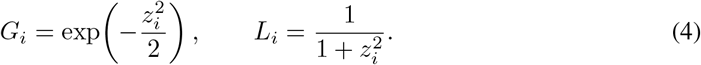

This ensemble-based weighting scheme derives from two intuitions: (a) a physically reasonable pose docked into the correct pocket should be relatively stable under small conformational perturbations – an outlier that disagrees with its local ensemble is unlikely to reflect the experimentally observed binding mode; and (2) ensemble consistency alone is insufficient – an absolute confidence score (e.g., the interface predicted TM-score, ipTM (Jumper et al., 2021), which measures the predicted quality of the protein-ligand interface) is needed to assess the structural quality of the binding interface. Therefore, multiplying the confidence scores by *G*_*i*_ or *L*_*i*_ simultaneously accounts for local ensemble agreement and absolute interface quality.

## 4 Experiments

### 4.1 Study Setup

We evaluate ACER on two tasks: (1) pocket exploration, and (2) ensemble-based ranking. Each stage is assessed on two datasets that directly challenge co-folding models on generalizability (see Appendix A for additional experimental details). Uncertainty is estimated from 1000 bootstrap iterations. All the results are reported in the mean and standard deviation for each metric and dataset.

#### 4.1.1 Datasets

**Allosteric binders** (Nittinger et al., 2025), comprising proteins present in the training set, which is dominated by orthosteric ligands, i.e., ligands binding to the primary active site rather than to the secondary allosteric sites that modulate protein activity indirectly. This dataset consists of 20 allosteric ligands across 17 proteins.

**Runs N’ Poses** (Škrinjar et al., 2025b), consisting of structures entirely unseen by the models. We filter systems released after Boltz-2’s training cutoff (1 June 2023) and focus on failure cases where top-ranked predictions of default co-folding models misplace the ligand (center-of-mass distance *>* 2 Å from the X-ray structure) – yielding 88 and 55 protein-ligand pairs for Boltz-2 and Protenix-v1, respectively. ACER, applied to each base model, is evaluated on these two subsets separately to assess model-agnostic benefit. Results are reported under two evaluation settings: (a) the full failure-case set, (b) a filtered set where only the cluster representatives of ligand systems (based on pocket and ligand shape similarity, Škrinjar et al. (2025b)) are considered, and 124 prevalent ligands are excluded (Škrinjar et al., 2025b), and (c) the full scope without restricting to wrong-pocket cases (*n* = 217) for benchmarking.

#### 4.1.2 Pocket Exploration

##### Experimental design

We apply the two pocket exploration strategies sequentially: (1) decoy ligand conditioning using five molecules (ligand plus four identical decoys) and five diffusion samples, followed by (2) pocket repulsion for five rounds. This setting guarantees the same sampling budget of 30 samples for each system. We then compare ACER extending Boltz-2 and Protenix-v1 against five baselines: (a) two ligand-free baselines based on volumetric or physicochemical properties, i.e., P2Rank (Krivák & Hoksza, 2018) and FPocket (Le Guilloux et al., 2009), and (b) three co-folding baselines: Boltz-2, OpenFold-3, and Protenix-v1.

##### Metrics

Success is measured by the Distance Center-to-Center (DCC) (Stepniewska-Dziubinska et al., 2020), *i.e*., the distance between the centers of mass (CoM) of the predicted and the ground-truth ligand (Appendix, Eq. A5). For P2Rank and FPocket – which do not co-fold complexes – DCC is measured between the predicted pocket center and the CoM of the ground-truth ligand. A pocket is considered correctly retrieved if the DCC falls below a given threshold (2Å or 3Å in this work).

#### 4.1.3 Ensemble-based Ranking

##### Experimental design

We apply ACER on Boltz-2 ^2^ and use the pair interface predicted TM-score (ipTM) (Jumper et al., 2021) as the baseline for ACER (*see* Appendix, Eq.A9) as a measure of the quality of the protein-ligand interface. We then follow a three-step ranking protocol: (1) We narrow the pocket candidate to *P* pockets to discard clearly unreliable interfaces. Using *P* = 10 or 20 can capture almost all the recoverable pockets (Appendix A, Table B4). (2) Within each pocket constraint, we generate *S* local conformational ensembles, resulting in *P* × *S* poses in total. (3) We rank the resulting poses by different criteria: (a) pair ipTM (used as a baseline), and (b) ensemble-normalized pair ipTM, using the Gaussian and Lorentzian weights (Eq. 4). Unless explicitly specified, we use *P* = 10 and *S* = 5. We additionally benchmark ACER against a conventional docking pipeline, i.e. P2Rank-to-co-folding baselin where P2Rank identifies the top-*N* candidate pockets, whose pocket residues served as binding constraints for a subsequent co-folding step.

##### Metrics

We measure ranking success by the pose accuracy of the top-*N* candidates sorted by each scoring metric, using ligand RMSD (L-RMSD) as the accuracy (Appendix, Eq. A6). The success rate is evaluated at 2 Å and 3 Å thresholds (as percentage of systems having an L-RMSD lower than the chosen threshold), combined with pose quality check byPoseBusters (Buttenschoen et al., 2024), ensuring that generated poses are not only geometrically close to the experimental structure but also physically plausible (*see* Appendix, A.3).

### 4.2 Results

#### 4.2.1 Pocket Exploration

##### Allosteric binders

Increasing the number of diffusion samples from the baseline models increases the likelihood of correct ligand placement into the right pocket (Table 1), suggesting that the underrepresented allosteric pockets are accessible but undersampled under default settings. ACER surpasses all baselines in terms of success rate (DCC lower than 2 Å or 3 Å, Table 1). Notably, ACER produces gains ranging from +11% to +13% on Boltz-2, and from +32% to +35% on Protenix-v1.

**Table 1:** Allosteric binding pocket identification. (*n* = 20). DCC success rate (%): fraction of systems where at least one predicted pocket falls within 2 Å or 3 Å of X-ray ligand center of mass.

| Method | DCC<2 Å<br>(%)↑ | DCC<3 Å<br>(%)↑ |
| --- | --- | --- |
| P2Rank | 25.0 ± 9.3 | 35.0 ± 10.9 |
| FPocket | 25.0 ± 9.9 | 55.0 ± 10.7 |
| OpenFold-3 (preview-2) | 35.0 ± 10.8 | 40.0 ± 10.9 |
| Boltz-2 (× 5 samples) | 40.0 ± 10.9 | 50.0 ± 10.9 |
| Boltz-2 (× 30 samples) | 55.0 ± 11.3 | 60.0 ± 10.9 |
| **ACER – Boltz-2** | **66.6 ± 9.6** | **73.5 ± 9.1** |
| Protenix-v1 (× 5 samples) | 25.0 ± 9.6 | 50.0 ± 11.4 |
| Protenix-v1 (× 30 samples) | 40.0 ± 10.8 | 50.0 ± 11.3 |
| **ACER – Protenix-v1** | **72.9 ± 9.5** | **75.8 ± 9.3** |

##### Failed-pocket subsets

ACER outperforms its respective base models (Table 2). ACER–Boltz-2 improves DCC success rate compared to Boltz-2 by +13.8% (2 Å) and +14.4% (3 Å) on the filtered subset (*n* = 46), and by +7.3% and +7.9% on the full set (*n* = 88), though all gains fall within the propagated uncertainty reflecting the limited sample size. Similarly, ACER–Protenix-v1 improves by +14.7% and +14.6% on the filtered subset (*n* = 30) and by +8.8% and +9.7% on the full set (*n* = 55) While not reaching statistical significance, the gains are consistent with the larger improvements observed on the allosteric dataset, suggesting a real but harder-to-detect effect on this more challenging subset. Finally, in the low-similarity bins (similarity ≤ 40%), co-folding models show pocket rescue rates near zero (Appendix B, Fig. D2), underperforming the ligand-free methods. With similarity up to 40%, ACER – Boltz-2 improves the DCC success rate of the baseline from 2.9% to 35.3% compared to the baseline (30 samples, Fig. D1).

**Table 2:** Pocket-failure recovery. ACER is evaluated in comparison with the wrongly identified pockets by each base model. ACER consistently correct places ligands into pockets by the base model, evaluated on the *Runs N’ Poses* wrong-pocket subset (Škrinjar et al., 2025b). DCC success rate (%) at 2 Å and 3 Å thresholds is reported for the full set and a filtered subset.

| Method | Boltz-2 failed-pocket subset |  |  |  | Protenix-v1 failed-pocket subset |  |  |  |
| --- | --- | --- | --- | --- | --- | --- | --- | --- |
| | Filtered ($n = 46$) | | Total ($n = 88$) | | Filtered ($n = 30$) | | Total ($n = 55$) | |
|  | DCC<2 Å<br>(%)↑ | DCC<3 Å<br>(%)↑ | DCC<2 Å<br>(%)↑ | DCC<3 Å<br>(%)↑ | DCC<2 Å<br>(%)↑ | DCC<3 Å<br>(%)↑ | DCC<2 Å<br>(%)↑ | DCC<3 Å<br>(%)↑ |
| P2Rank | 24.9 ± 8.9 | 45.8 ± 10.2 | 22.9 ± 6.2 | 43.1 ± 7.3 | 20.9 ± 9.1 | 42.9 ± 11.2 | 23.8 ± 7.6 | **49.7 ± 9.4** |
| FPocket | 19.5 ± 5.9 | 37.0 ± 7.0 | 27.2 ± 4.8 | 42.1 ± 5.3 | 10.2 ± 5.6 | 36.7 ± 8.7 | 22.1 ± 5.8 | 40.8 ± 6.5 |
| Boltz-2 (× 30 samples) | 30.7 ± 6.6 | 37.3 ± 7.1 | 36.1 ± 5.2 | 42.9 ± 5.3 | – | – | – | – |
| **ACER – Boltz-2** | **44.5 ± 7.1** | **51.7 ± 7.0** | **43.4 ± 5.0** | **50.8 ± 5.0** | – | – | – | – |
| Protenix-v1 (× 30 samples) | – | – | – | – | 19.4 ± 6.9 | 30.1 ± 8.0 | 19.2 ± 5.1 | 27.7 ± 5.7 |
| **ACER – Protenix-v1** | – | – | – | – | **34.1 ± 7.9** | **44.7 ± 7.8** | **28.0 ± 5.8** | 37.4 ± 5.9 |

#### 4.2.2 Ensemble-based Ranking

##### Allosteric binders

ACER – L-weight consistently performs on par with or outperforms all baselines. At Top-5, ACER Boltz-2 – L-weight gains +20.0% and +25.0% at 2 Å and +25.0% at 3 Å over the strongest baseline (Boltz-2). At Top-1, ACER matches Boltz-2. Unlike pair ipTM from static snapshots (Table 3), the L-weighted and G-weighted variants leverage local conformational ensembles, thereby recovering accurate poses that static-pose ranking would otherwise miss.

**Table 3:**
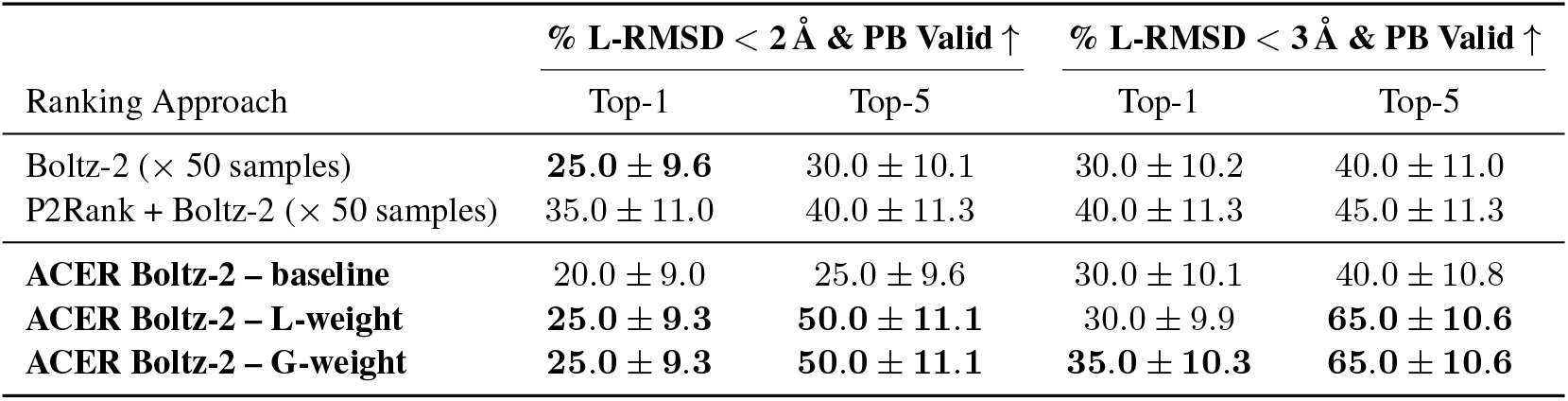
Pose ranking on the allosteric set. Top-*k* L-RMSD (%) at 2 Å and 3 Å thresholds is reported, considering only physically valid poses (Buttenschoen et al., 2024). *ACER Boltz-2 – baseline* uses raw pair ipTM as scoring function; *ACER Boltz-2 – L-weight* and *ACER Boltz-2 – G-weight* weight pair ipTM by the Lorentzian and Gaussian (Eq. 4) ensemble normalization, respectively.

##### Runs N’ Poses

On the failed-pocket subset of the *Runs N’ Poses* benchmark (Table 4), ACER consistently outperforms the Boltz-2 baseline across both thresholds (gains at +8.5% and +4.6% on the filtered and full sets, respectively). ACER achieved the largest gains at Top-5 on the filtered subset (23.9% → 28.3% at 2 Å and 32.6% → 39.1% at 3 Å).

**Table 4:** Pose ranking on the *Runs N’ Poses* wrong-pocket subset of Boltz-2. Top-*k* L-RMSD (%) at 2 Å and 3 Å thresholds is reported, considering only physically valid poses (Buttenschoen et al., 2024)). *ACER – baseline* uses raw pair ipTM as scoring function; *ACER – L-weight* and *ACER – G-weight* weight pair ipTM by the Lorentzian and Gaussian (Eq. 4) ensemble normalization.

| Ranking Approach | Clustered & Distinct Ligands ($n = 46$) | | | | Total ($n = 88$) | | | |
| --- | --- | --- | --- | --- | --- | --- | --- | --- |
| | % L-RMSD < 2 Å & PB Valid ↑ | | % L-RMSD < 3 Å & PB Valid ↑ | | % L-RMSD < 2 Å & PB Valid ↑ | | % L-RMSD < 3 Å & PB Valid ↑ | |
|  | Top-1 | Top-5 | Top-1 | Top-5 | Top-1 | Top-5 | Top-1 | Top-5 |
| Boltz-2 (× 50 samples) | 15.2 ± 5.5 | 15.2 ± 5.5 | 19.6 ± 6.0 | 19.6 ± 6.0 | 26.1 ± 4.8 | 27.3 ± 4.8 | 28.4 ± 4.9 | 29.5 ± 5.0 |
| **ACER Boltz-2 – baseline** | **23.9 ± 6.4** | 23.9 ± 6.4 | **32.6 ± 7.1** | 34.8 ± 7.2 | **30.7 ± 5.0** | 31.8 ± 5.1 | **35.2 ± 5.2** | 37.5 ± 5.3 |
| **ACER Boltz-2 – L-weight** | 15.2 ± 5.5 | 26.1 ± 6.7 | 21.7 ± 6.3 | 37.0 ± 7.2 | 21.6 ± 4.4 | 29.5 ± 4.9 | 26.1 ± 4.8 | 36.4 ± 5.3 |
| **ACER Boltz-2 – G-weight** | 15.2 ± 5.5 | **28.3 ± 6.8** | 21.7 ± 6.3 | **39.1 ± 7.2** | 21.6 ± 4.4 | **31.8 ± 5.1** | 26.1 ± 4.8 | **38.6 ± 5.4** |

To benchmark ACER against all co-folding baselines, we extend the analysis to the full *Runs N’ Poses* set (*n* = 217, clustered and distinct ligands, Fig. 2), without restricting it to wrong-pocket cases. On this set, ACER achieves the highest Top-5 success rate (52.5%) among all co-folding models, outperforming Boltz-2 (49.8%), Protenix-v1 (51.2%), and OpenFold-3 (46.1%). At Top-1, ACER outperforms Protenix-v1 and OpenFold-3 but falls behind Boltz-2 (47.0% → 40.0%), suggesting that ensemble-based scoring improves candidate diversity but might struggle to promote the best pose to the very top position. Notably, ACER is the only method to achieve a non-zero top-1 success rate (7%) in the lowest-similarity bin (up to 20% similarity) (Fig. 2a). When increasing the sampling budget to 200 samples (*P* = 20 and *S* = 10), ACER successfully samples accurate poses, reaching 21% at Top-20 and 28% at 200-oracle, demonstrating that its generalization can be improved with additional pocket candidates and higher coverage from local ensembles (Fig. 2b,c).

**Figure 2:**
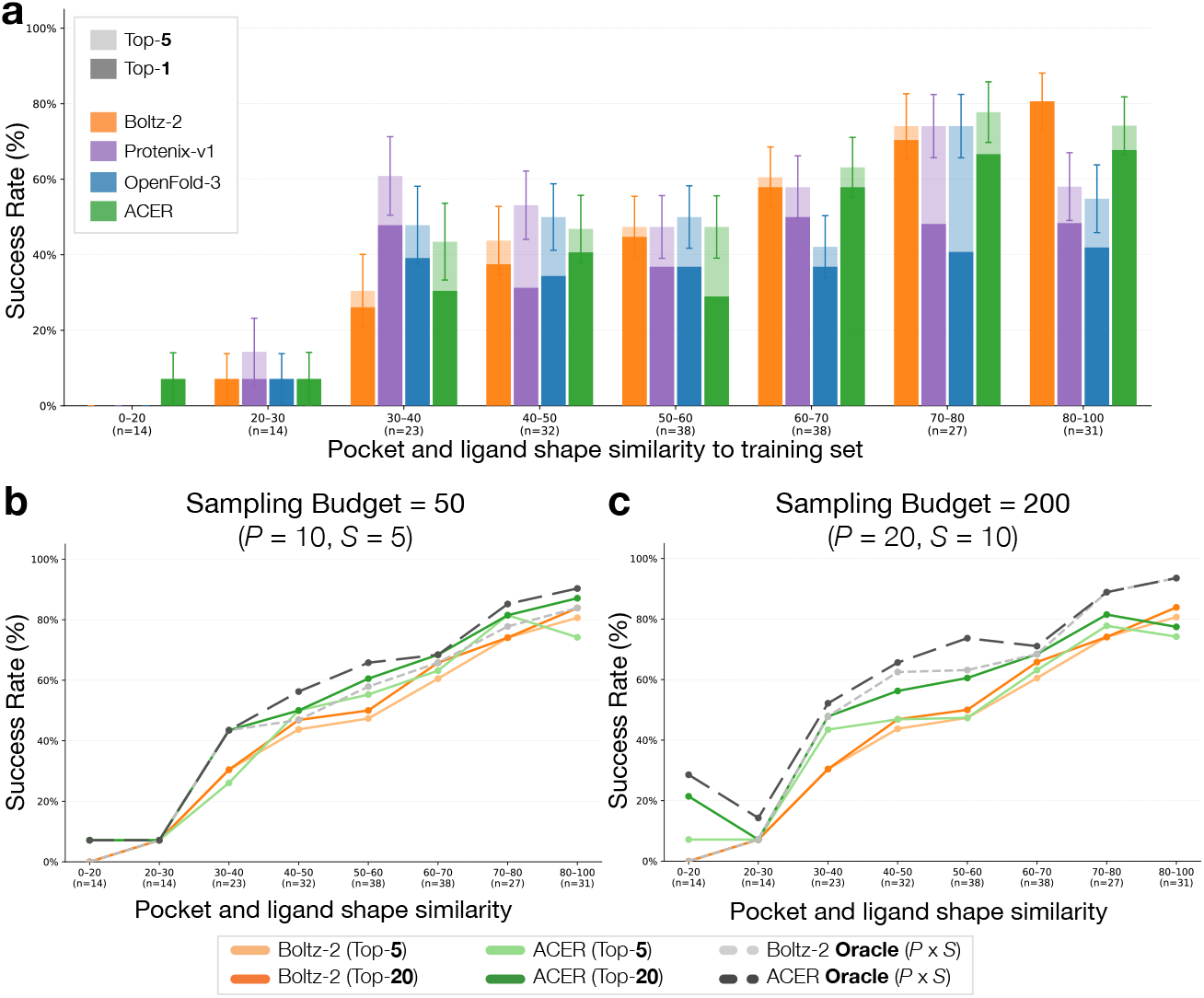
Ranking on *Runs N’ Poses*, measured as success rate (percentage of L-RMSD *<* 2 Å and PB valid). **(a)** ACER vs co-folding baselines on the clustered and distinct ligand subset (*n* = 217). **(b, c)** Success rate of Boltz-2 and ACER at sampling budgets of **(b)** 50 and **(c)** 200 samples. Increasing the budget rescues accurate poses in the hardest similarity bin (0–20] particularly at top-20. (*P* = no. of pocket candidates to consider; *S* = no. of local conformation ensembles).

##### Failure modes

To better understand ACER’s benefits, we further analyze all the poses per system from local ensembles from ACER Boltz-2 – L-weighted for the Boltz-2 failed-pocket subset. We identify three main failure modes (Fig. D4): (1) *Correct pocket, low rank* (14.7% of cases). ACER successfully places ligands into the correct pockets but fails to rank them highly to top 5 candidates, indicating the limitation of the ranking procedure (Fig. D4a and Fig. D1). (2) *Correct ligand placement but wrong pose* (26.5% of cases). The ligand placement is correct, but an accurate pose is never generated; this is likely due to ineffective modeling of difficult protein–ligand interfaces by the baseline co-folding model (Fig. D4b). (3) *Correct pocket not rescued* (58.8% of cases), leaving room for further improvement in pocket exploration (Fig. D4c).

#### 4.2.3. Runtime performance

ACER incurs additional compute overhead compared to the baselines: each decoy ligand in decoy ligand conditioning adds ~2× wall-clock time (trajectories are conditioned on reconstructed cached representation) (Table C6 and C7 in Appendix), and iterative pocket repulsion introduces ~3× overhead per round of guided sampling due to SMC resampling. Together, both approaches accumulate to ~5× per diffusion sample in total. Ranking overhead is modest and comparable to the baseline. Despite this cost, ACER successfully rescues binding pockets and poses that are undersampled under default co-folding. ACER is hence positioned as a *precision-oriented tool* for challenging targets – such as allosteric or structurally novel systems – where accurate binding site characterization justifies the overhead.

## 5 Limitations

While ACER demonstrates consistent improvements on challenging targets, several avenues for improvement remain. The ~5× runtime overhead currently limits applicability in high-throughput settings, though this could be reduced through more efficient sampling strategies or caching mechanisms. Pocket recovery on *Runs N’ Poses* reaches ~50% within a budget of 30 samples – a promising result on hard targets, but one that leaves clear room for improvement as sampling budgets and exploration strategies mature. Ensemble-based ranking consistently improves top-5 and beyond, and closing the gap to top-1 represents a natural next step, likely addressable through better scoring functions or learned ranking models. Finally, statistical validation remains challenging given the scarcity of allosteric and out-of-distribution targets – a limitation shared across the field, and one that underscores the need for larger, more diverse benchmarks for generalizable co-folding. Further evaluation on the recent dataset (OpenBind Consortium, 2026) would be a valuable next step.

## 6 Discussion

Co-folding models have emerged as a tool for predicting protein-ligand structures, yet their tendency to memorize the training set distributions can lead to systematic ligand misplacement on allosteric binding and out-of-distribution targets – precisely the most relevant cases for drug discovery. ACER addresses this through a training-free framework combining decoy ligand conditioning and iterative pocket repulsion to redirect the model’s sampling toward underrepresented binding sites. Its ensemble-based ranking further enables a principled comparison of binding poses across candidate pockets, providing an informative starting point for downstream refinement, such as energy minimization, specialized docking (Prat et al., 2026), or extensive MD simulations. Taken together, these results suggest that the generalizability gap in co-folding is not a fundamental limitation of the underlying models, but rather a sampling problem amenable to inference-time intervention. ACER demonstrates that redirecting the denoising trajectory – without touching model weights – can unlock binding sites that are systematically missed under default sampling. This reframing opens a broader research direction: as co-folding models continue to improve, inference-time exploration strategies like ACER may prove essential for extending their reach to the underrepresented pocketome. This suggests a further application to fragment-based virtual screening, where ACER could be used to co-fold small fragments as probes to identify alternative binding pockets and characterize their interaction patterns. A recent study (Purnomo et al., 2026) explores this related fragment-screening strategy for identifying allosteric and cryptic binding sites in kinase proteins, with findings that are consistent with the mechanistic insights provided by ACER.

Looking beyond protein-ligand interactions, extending ACER’s rationale to protein-protein or protein-nucleic acid interfaces (where co-folding models are similarly susceptible to memorization) represents a natural and important future direction. ACER establishes a blueprint for training-free generalization in structural biology: rather than collecting more data or retraining larger models, one can instead design smarter sampling dynamics that push existing models beyond their training distribution.

## Impact statement

This work advances computational methods for protein-ligand structure prediction, with direct applications to structure-based drug discovery. By enabling co-folding models to explore the underrepresented pocketome, ACER accelerates the identification of novel therapeutic targets and pharmaceutical drugs that are currently inaccessible to standard computational pipelines.

## Appendix

### A. Experimental Details

#### A.1 Dataset Details

##### Allosteric systems

We adopt the same set of allosteric complexes from Nittinger et al. (2025) where the details are provided in their supplementary materials. The dataset was originally published by Xie et al. (2022) and Olanders et al. (2024). To enhance the quality of the structures, the structures are removed if they are (1) close analogues binding to both orthosteric and allosteric binding site, (2) with different UniProt Ids, (3) where ligands bind to different proteins, and (4) with covalent ligands or peptides. The details of individual system is provided in Table A1.

**Table A1:**
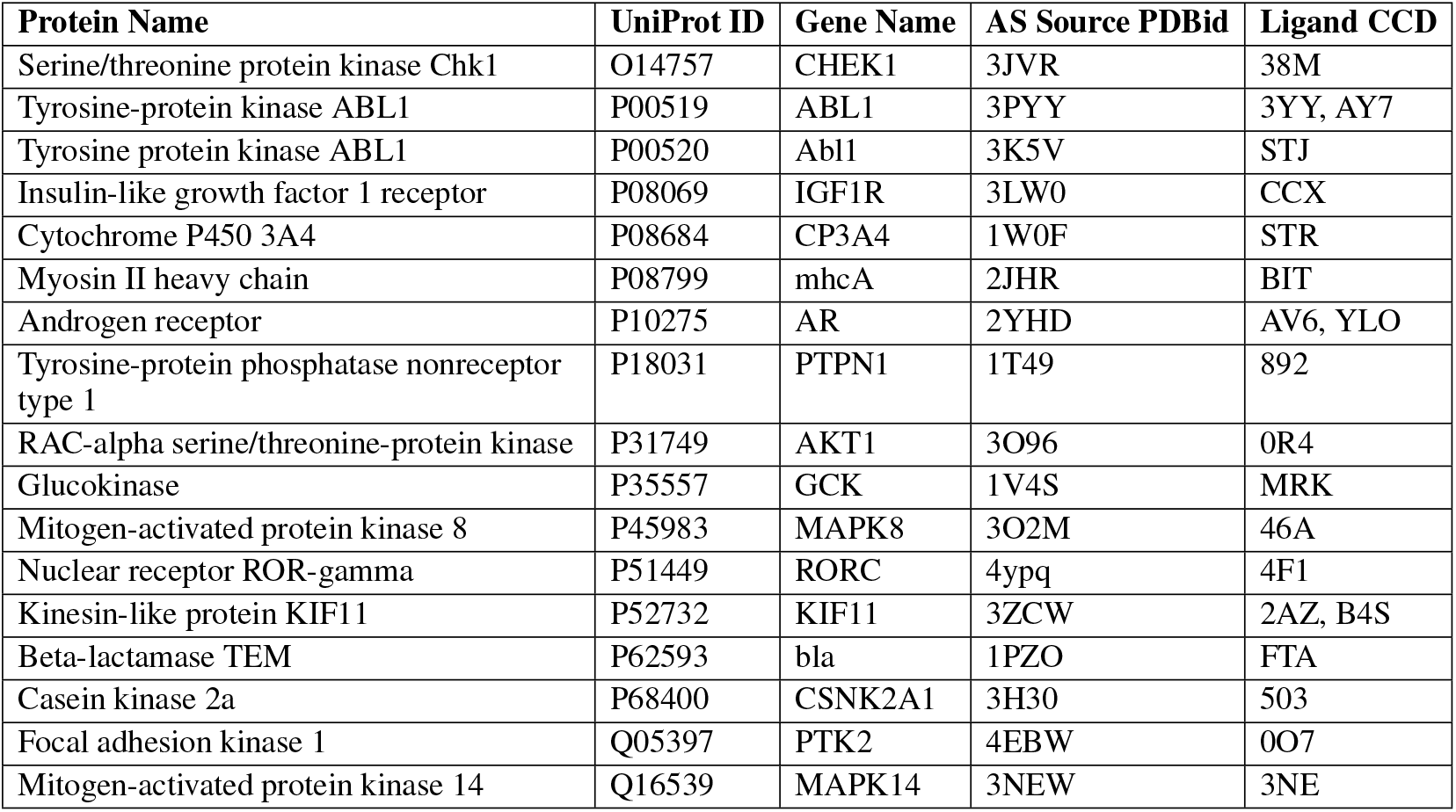
Protein targets with UniProt IDs, gene names, PDB IDs, and ligand CCDs for 20 allosteric compounds.

| Protein Name | UniProt ID | Gene Name | AS Source PDBid | Ligand CCD |
| --- | --- | --- | --- | --- |
| Serine/threonine protein kinase Chk1 | O14757 | CHEK1 | 3JVR | 38M |
| Tyrosine-protein kinase ABL1 | P00519 | ABL1 | 3PYY | 3YY, AY7 |
| Tyrosine protein kinase ABL1 | P00520 | Abl1 | 3K5V | STJ |
| Insulin-like growth factor 1 receptor | P08069 | IGF1R | 3LW0 | CCX |
| Cytochrome P450 3A4 | P08684 | CP3A4 | 1W0F | STR |
| Myosin II heavy chain | P08799 | mhcA | 2JHR | BIT |
| Androgen receptor | P10275 | AR | 2YHD | AV6, YLO |
| Tyrosine-protein phosphatase nonreceptor type 1 | P18031 | PTPN1 | 1T49 | 892 |
| RAC-alpha serine/threonine-protein kinase | P31749 | AKT1 | 3O96 | 0R4 |
| Glucokinase | P35557 | GCK | 1V4S | MRK |
| Mitogen-activated protein kinase 8 | P45983 | MAPK8 | 3O2M | 46A |
| Nuclear receptor ROR-gamma | P51449 | RORC | 4ypq | 4F1 |
| Kinesin-like protein KIF11 | P52732 | KIF11 | 3ZCW | 2AZ, B4S |
| Beta-lactamase TEM | P62593 | bla | 1PZO | FTA |
| Casein kinase 2a | P68400 | CSNK2A1 | 3H30 | 503 |
| Focal adhesion kinase 1 | Q05397 | PTK2 | 4EBW | 0O7 |
| Mitogen-activated protein kinase 14 | Q16539 | MAPK14 | 3NEW | 3NE |

##### Runs N’ Poses

Runs N’ Poses is a well-established benchmark for protein–ligand generalization that groups samples by pocket similarity, pocket sequence identity, ligand similarity, and shape overlay (Škrinjar et al., 2025b). Because we apply ACER on top of Boltz-2, we restrict evaluation to systems deposited after its training cutoff of 1 June 2023. To isolate the wrong-pocket failure mode that ACER is designed to address, we further filter to systems where the baseline prediction misplaces the ligand, defined as a center-of-mass distance greater than 2 Å from the X-ray reference structure. The protein-ion pairs are excluded from the evaluation. The resulting test set is stratified by closest similarity to pre-cutoff training samples, allowing us to analyze performance as a function of target difficulty. All evaluations are reported on two subsets: (i) the full set of post-cutoff systems, and (ii) a clustered and filtered variant that removes 124 prevalent ligands as in Škrinjar et al. (2025b), retaining only non-overlapping systems to reflect genuine gains in generalization.

To establish the protein-ligand generalizability benchmark, we extend our evaluation to the full scope of Runs N’ Poses. We remain focused on the clustered and distinct ligands subset, as it represents the most challenging subset for generalization and avoids double-counting successes on similar structures. Following the protocol prescribed by Škrinjar et al. (2025b) and running the code from their official repository, we obtain a final set of 217 systems released after 1 June 2023.

#### A.2 Parameter Settings

##### Co-folding model settings

We report Boltz-2’s sampling hyperparameters used across our experiments in Table A2.

**Table A2:** Boltz-2 sampling hyperparameters Hyperparameter.

| Hyperparameter | Value |
| --- | --- |
| num_particles | 3 |
| fk_lambda | 4 |
| fk_resampling_interval | 3 |
| inner_gradient_iterations | 20 |

##### Pocket exploration

ACER is implemented on top of Boltz-2 and Protenix-v1, which we select because it is open-source and supports both inference-time steering and local ensemble generation. Exploration proceeds in two phases applied in sequence. In the first phase, we run the inference of 5 diffusion samples with different number of decoy ligands (1 through 5), where each decoy is a duplicate of the target ligand. In the second phase, we perform 5 additional inference iterations with pocket repulsion, using *c*_*g*_ computed from pocket residues within 6 Å of ligand atoms, *d*_min_ = 5 Å, and *r*_*g*_ = 8 Å. We report the mean and standard deviation of DCC success rate at which uncertainty estimates are computed over 5 seeds (seeds = 231234, 89567, 412903, 451027, 738291). We ablate the parameters associated with each method in detail in D

##### Ensemble-based ranking

For ranking tasks, we take the output from a single seed of the exploration stage and pocket constraints from residues whose atoms lie within 6 Å of any ligand atom from top *P* pockets ranked by pair ipTM. Leveraging those pocket constraints, We run the inference with *S* diffusion samples with MD method conditioning to generate local conformational ensembles. The total sampling budget is therefore *P* × *S* samples. Unless explicitly specified, we use *P* = 10 and *S* = 5 in the main experiment. Unlike pocket exploration, we only report the results and corresponding uncertainty estimates on a single seed (seed = 738291). The ablation study on the total sampling budget is provided in Section D.

#### A.3 Metrics

##### Pocket recovery

For multi-chain systems that contain duplicated protein-ligand pairs, we first superpose the protein chain of predicted structure onto the ground-truth chain by backbone alignment. Similarly to Ma et al. (2025), we handle homodimers or copies of equivalent protein chains in the reference structure by permuting the alignment between predicted and ground-truth chains to obtain the best results, as formulated below.

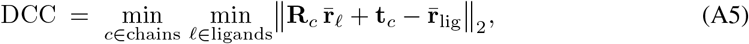

where (**R**_*c*_, **t**_*c*_) is the rigid-body transform from the prediction frame to ground-truth chain *c* and 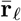 is the heavy-atom centroid of predicted ligand chain ℓ (waters and hydrogens excluded). Minimum over *c* and ℓ is the permutation-invariant best alignment; a system is counted as *rescued* at threshold *τ* if at least one predicted ligand has DCC *< τ*.

##### Pose accuracy

Per-pose ligand RMSD (L-RMSD) is computed as:

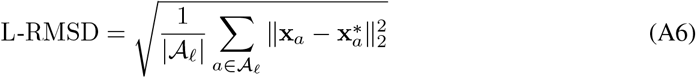

where *A*_ℓ_ denotes the set of ligand heavy atoms, **x**_*a*_ ∈ ℝ^3^ is the predicted position of atom *a*, and 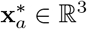 is its ground-truth position. L-RMSD is computed with OpenStructure (Biasini et al., 2013). The software first establishes a chain correspondence between prediction and reference using its lDDT-based chain-mapping algorithm. Symmetry-aware atom correspondences are enumerated via molecular-graph automorphism on the heavy-atom skeleton and the reported *symmetry-corrected ligand RMSD* is the minimum over all valid atom mappings after superposing the backbone carbon of the assigned protein chain. This yields a permutation-invariant pose-quality score, robust both to protein-chain renaming and to ligand internal symmetry.

##### Pose quality

Besides structural accuracy, we assess whether poses are physically plausible. We use PoseBusters plausibility checks using the PoseBusters package version 0.6.5. The pose is considered valid only if it passes all the following checks.

|  |  |
| --- | --- |
| `mol_pred_loaded` | `internal_steric_clash` |
| `mol_true_loaded` | `aromatic_ring_flatness` |
| `mol_cond_loaded` | `double_bond_flatness` |
| `sanitization` | `internal_energy` |
| `molecular_formula` | `minimum_distance_to_protein` |
| `molecular_bonds` | `minimum_distance_to_organic_cofactors` |
| `double_bond_stereochemistry` | `minimum_distance_to_inorganic_cofactors` |
| `tetrahedral_chirality` | `volume_overlap_with_protein` |
| `bond_lengths` | `volume_overlap_with_organic_cofactors` |
| `bond_angles` | `volume_overlap_with_inorganic_cofactors` |

##### Pair ipTM

ipTM score, introduced in (Jumper et al., 2021), quantifies the alignment quality between two molecular entities *A* and *B*. It is defined asymmetrically as:

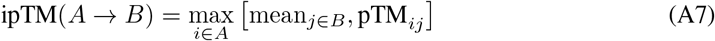

where the predicted TM score (pTM) computes the TM-score from predicted aligned errors (PAE):

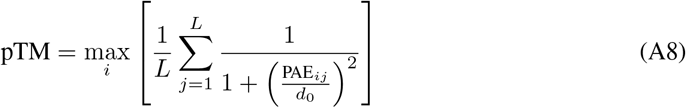

Jumper et al. (2021); Abramson et al. (2024) provides ipTM for a pair of molecular entities. In protein-ligand complexes, it provides 2 values of directional ipTM. Therefore, we define the pair ipTM as:

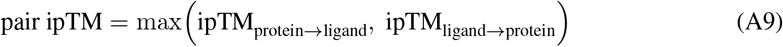

#### A.4 Baseline Details

##### Pocket exploration

To establish baselines for pocket exploration, we consider both traditional pocket detection methods and modern co-folding approaches. For the former, P2Rank (Krivák & Hoksza, 2018) predicts ligand binding sites using a random forest classifier trained on local physicochemical features of the protein surface. FPocket (Le Guilloux et al., 2009) detects cavities on the protein surface using Voronoi tessellation, without requiring any training data or ligand information. Both produce ranked lists of candidate pocket centers without explicit ligand placement. For these methods, DCC is measured between the predicted pocket center and the ground-truth ligand center of mass. For co-folding baselines, we use the same inference protocols as ACER for Boltz-2, Protenix-v1 (protenix_base_default_v1.0.0), and OpenFold-3 (preview-2) to generate 30 samples but without decoy ligand conditioning or pocket repulsion potential. For Boltz-2 and Protenix-v1, we also enable the default potentials to ensure physical plausibility of the poses.

##### Ensemble-based ranking

To benchmark the pose ranking scheme, we generate *P × S* samples (*P* = 10, *S* = 5) using Boltz-2, Protenix-v1, and OpenFold-3, ranked by each model’s confidence scores, maintaining equal sampling budget across all methods for a fair comparison with ACER. When reporting best-of-k or top-k results in tables and figures, we consider the best pose among top k candidates, which is both physically valid and has the lowest L-RMSD within the threshold.

### B Extended Results

#### B.1 Mechanism of decoy ligand conditioning

To illuminate the mechanism of decoy ligand conditioning, we analyse how trunk representations and denoiser attention weights change on the 9 allosteric systems that Boltz-2 misses, with and without decoy ligand conditioning (Table B1).

##### Distogram shifts in trunk representations

The distogram shows a higher preference toward allosteric pocket residues (tokens within 4 Å of any ligand atom). Let 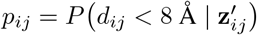 be the distogram-predicted contact probability conditioned on 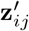. We aggregate over a pocket residue set *ℛ* and the atoms *ℒ* of the queried ligand ℓ as

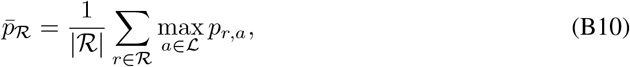

the mean strongest predicted contact per ligand. This measures the geometric bias that the trunk imposes on the denoiser.

**Table B1:** Trunk and denoiser statistics on the 9 allosteric systems that Boltz-2 misses, averaged over systems. Ratios above 1 indicate a bias toward the allosteric rather than the orthosteric pocket.

| Metric | Without decoys | With decoys |
| --- | --- | --- |
| *Trunk (distogram contact preference)* |  |  |
| $\bar{p}_{\mathcal{R}_{\text{allo}}}$ | 0.045 | **0.216** |
| $\bar{p}_{\mathcal{R}_{\text{ortho}}}$ | 0.551 | 0.380 |
| *Denoiser (allosteric-to-orthosteric ratios)* |  |  |
| $\ \mathbf{z}'\ _{\text{allo}}/\ \mathbf{z}'\ _{\text{ortho}}$ | 0.82 | **1.09** |
| $\mathbb{E}_{\text{allo}}[w_{ij}^{\text{pre}}]/\mathbb{E}_{\text{ortho}}[w_{ij}^{\text{pre}}]$ | 0.381 | **1.931** |
| $\mathbb{E}_{\text{allo}}[w_{ij}^{\text{post}}]/\mathbb{E}_{\text{ortho}}[w_{ij}^{\text{post}}]$ | 0.494 | **1.598** |

##### Shifts in denoiser’s attention weights

Conditioned on decoys, the attention weights of the token_transformer on alternative pocket residues increase. These weights are shaped by two components: the raw query–key attention and the pair-representation bias. To quantify the effect of the decoy ligand on each, we compute the pre-bias attention weight

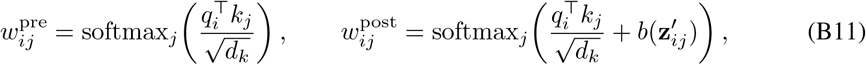

where 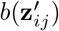 is the learned pair-bias term, together with the norm 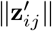, for *i, j* ranging over ligand atoms and pocket residues.

Average contact probability shifts toward the allosteric pocket (0.045 → 0.216), while the orthosteric value falls from 0.551 to 0.380. The allosteric-to-orthosteric ratios of the pre- and post-bias attention weights both rise above 1, indicating a stronger geometric bias toward allosteric residues. Although the pre-bias weights exclude 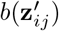, the altered representations already propagate into the denoiser through diffusion conditioning of the token embeddings. Decoy ligands thus occlude the dominant pocket, mitigating memorisation and allowing greater attention on alternative sites.

#### B.2 Characteristics of ACER’s explored pockets

Reported performance (Table 1-2) measures the recovery of known binding pockets considered biologically relevant. Table B2 shows that ACER also rescues *unfamiliar* pockets that are structurally and chemically dissimilar to the training set, i.e. those falling in the 0–20 similarity bin. Recovery alone, however, does not capture whether a method is useful for discovering *alternative* druggable sites. We therefore characterise the diversity and novelty of the pockets explored by ACER and by P2Rank on the 20-system allosteric set.

##### Diversity

We analyse inter-pocket-centre distances and pocket radius of gyration to quantify how pockets disperse over the protein surface. Distinct pockets are obtained by union–find clustering at 2 Å: two pockets are merged into the same cluster if their centres lie within 2 Å. Pocket centres are taken as the ligand centre of mass for ACER, and as the P2Rank-reported pocket centre for P2Rank.

##### Novelty

We characterise pockets structurally and physicochemically against Boltz-2’s default pocket. Structural novelty is measured by the pocket shape similarity of *Runs N’ Poses* (Škrinjar et al., 2025b) and by the pocket-aligned TM-score; physicochemical novelty by the Jensen–Shannon (JS) divergence between histograms of non-covalent interactions between pocket residues and ligand atoms, computed with ProLIF. Lower shape similarity and TM-score, and higher JS divergence, indicate a pocket further from the default site.

**Table B2:** Diversity and novelty of explored pockets on the allosteric set (*n* = 20), averaged over systems. Novelty metrics are computed against Boltz-2’s default pocket and are therefore undefined for P2Rank, which does not produce a ligand pose.

| Method | Diversity |  |  | Novelty |  |  |
| --- | --- | --- | --- | --- | --- | --- |
|  | Distinct pockets | Inter-pocket distance (Å) | Radius of gyration (Å) | Shape similarity ↓ | JS ProLIF ↑ | Pocket-aligned TM-score ↓ |
| ACER – Boltz-2 | 13.8 | 24.1 | 7.9 | 0.085 | 0.30 | 0.63 |
| P2Rank | 8.2 | 32.7 | 7.4 | 0.040 | – | – |

**Table B3:**
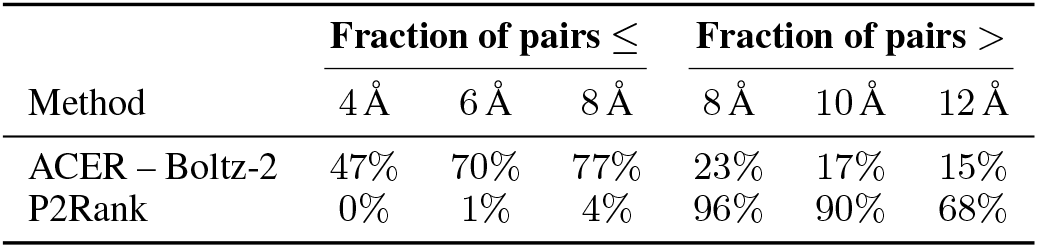
Cumulative distribution of inter-pocket-centre distances. Each entry is the fraction of distinct-pocket pairs whose centres are separated by less than (left block) or more than (right block) the stated distance.

ACER proposes roughly 1.7× more distinct pockets per system than P2Rank (13.8 vs. 8.2) at a finer spatial granularity: its pocket centres are closer together on average (24.1 Å vs. 32.7 Å), and 47% of pocket pairs lie within 4 Å of one another, against 0% for P2Rank (Table B3). This addresses the concern that repulsion steering could be risky when the model needs to refine around a nearby correct site, or when several valid sites lie close together: ACER retains the resolution to separate such sites rather than pushing sampling away from the whole region. The granularity is also tunable through the pocket radius used in repulsion steering.

At the same time, ACER’s exploration is not confined to a small patch of the protein surface, as 23% of pocket pairs remain more than 8 Å apart. P2Rank, by contrast, proposes a smaller number of more widely dispersed pockets, so the two tools are complementary and offer control over how localised the exploration should be. The novelty metrics (Table B2) show that ACER’s pockets depart from Boltz-2’s default site both in shape and in interaction profile, indicating that ACER can propose cryptic pockets with distinct shape and chemical complementarity to fragment probes, which suits it to fragment-based discovery.

##### Top pocket candidate selection

We show how %DCC success rate varies by the top *P* pockets according to pair ipTM to discard clearly unreliable predicted interfaces. Table B4 shows how selected top *P* pockets captures recoverable pockets across subsets, achieving 100% rescue rates at Top-20 under both DCC criteria, while avoiding running local ensemble generation for all the pocket candidates, which can be computationally expensive in the subsequent ranking stage.

**Table B4:**
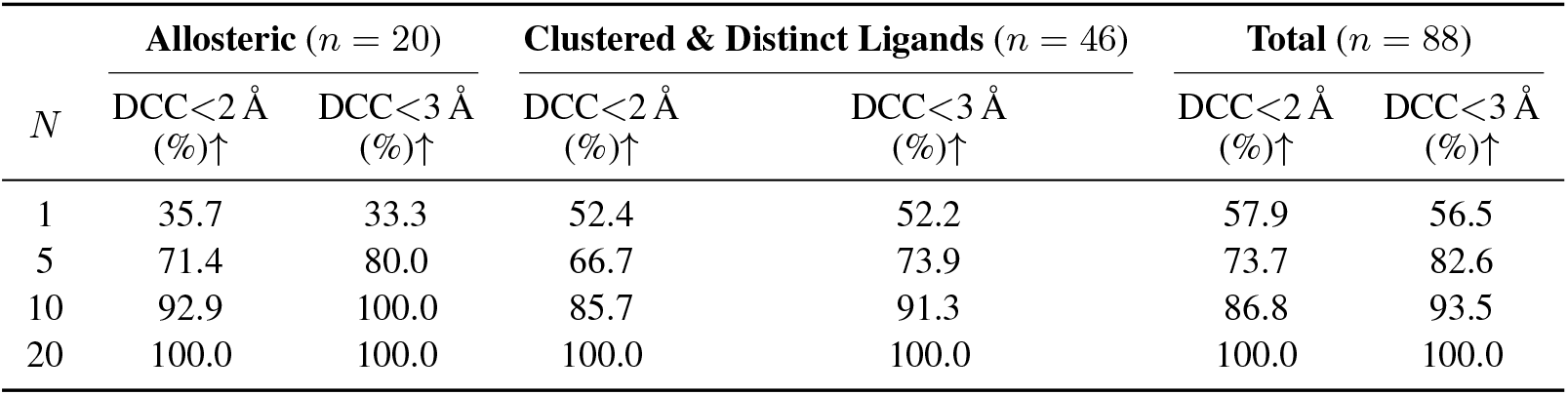
Pocket rescue rate for the top-*N* pockets ranked by pocket exploration strategies. DCC *<* 2 Å and *<* 3 Å denote the distance criteria for pocket recovery. Percentages are computed relative to the ceiling of recoverable pockets.

| $N$ | Allosteric ($n = 20$) | | Clustered & Distinct Ligands ($n = 46$) | | Total ($n = 88$) | |
| --- | --- | --- | --- | --- | --- | --- |
| | DCC $< 2$ Å (%)↑ | DCC $< 3$ Å (%)↑ | DCC $< 2$ Å (%)↑ | DCC $< 3$ Å (%)↑ | DCC $< 2$ Å (%)↑ | DCC $< 3$ Å (%)↑ |
| 1 | 35.7 | 33.3 | 52.4 | 52.2 | 57.9 | 56.5 |
| 5 | 71.4 | 80.0 | 66.7 | 73.9 | 73.7 | 82.6 |
| 10 | 92.9 | 100.0 | 85.7 | 91.3 | 86.8 | 93.5 |
| 20 | 100.0 | 100.0 | 100.0 | 100.0 | 100.0 | 100.0 |

##### Multi-seed results

To assess whether wrong-pocket predictions are consistent across random seeds, we repeat each inference 5 times and compute the union of successful placements. Concretely, a system is counted as successful if at least one of the 5 predictions achieves a DCC below the threshold, regardless of the other seeds. As shown in Figure D3, increasing the number of samples modestly reduces the fraction of wrong-pocket systems compared to the single top-1 prediction.

##### Pocket and pose recovery by similarity bins

Figure D1 shows the best L-RMSD per system stratified by pocket-ligand shape similarity to the training set, restricted to systems where the Boltz-2 baseline fails to identify the correct pocket (DCC ≥ 2 Å). ACER rescues are most prevalent in the low-similarity regime (0–30), where Boltz-2 consistently misplaces ligands, and in several of these rescued cases ACER can achieve an accurate pose (L-RMSD *<* 2 Å), despite the ranking bottleneck, as described in 4.2.2.

##### Protein-ligand generalizability benchmark

Table B5 details the success rate (% L-RMSD < 2 Å or 3 Å thresholds and PB-Valid). While ACER’s top-1 performance is inferior to co-folding baselines, it offers clear advantages at Top-5 and Top-10 success rate at which. it can sample accurate poses in the challenging similarity bins (0 –30]. On the overall benchmark, ACER achieves 52.5% and 54.8% at Top-5 and Top-10 (L-RMSD < 2 Å) with a similar trend at the 3 Å threshold (64.1% and 65.9%). This pattern indicates that ACER’s ranking effectively high-quality poses within a small candidate pool, even though its top-ranked pose does not lead.

**Table B5:** Performance comparison of co-folding methods across sequence identity bins on the Runs N’ Poses benchmark (*n* = 217 when top *P* = 10 pocket candidates and *S* = 5 samples of local ensembles are considered).

| Similarity bin (n) | Ranking Approach | % L-RMSD < 2 Å & PB Valid ↑ |  |  | % L-RMSD < 3 Å & PB Valid ↑ |  |  |
| --- | --- | --- | --- | --- | --- | --- | --- |
|  |  | Top-1 | Top-5 | Top-10 | Top-1 | Top-5 | Top-10 |
| 0–20 (14) | Boltz-2 | 0.0 ± 0.0 | 0.0 ± 0.0 | 0.0 ± 0.0 | 0.0 ± 0.0 | 0.0 ± 0.0 | 0.0 ± 0.0 |
|  | Protenix-v1 | 0.0 ± 0.0 | 0.0 ± 0.0 | 0.0 ± 0.0 | 0.0 ± 0.0 | 0.0 ± 0.0 | 0.0 ± 0.0 |
|  | OpenFold-3 | 0.0 ± 0.0 | 0.0 ± 0.0 | 7.1 ± 6.8 | 0.0 ± 0.0 | 7.1 ± 6.8 | 7.1 ± 6.8 |
|  | ACER Boltz-2 L-weight | **7.1 ± 7.1** | **7.1 ± 7.1** | **7.1 ± 7.1** | **7.1 ± 7.1** | **7.1 ± 7.1** | **14.3 ± 9.3** |
| 20–30 (14) | Boltz-2 | 7.1 ± 6.8 | 7.1 ± 6.8 | 7.1 ± 6.8 | 7.1 ± 6.8 | 7.1 ± 6.8 | 7.1 ± 6.8 |
|  | Protenix-v1 | 7.1 ± 6.8 | **14.3 ± 9.4** | **14.3 ± 9.4** | 7.1 ± 6.8 | **14.3 ± 9.4** | **14.3 ± 9.4** |
|  | OpenFold-3 | 7.1 ± 6.8 | 7.1 ± 6.8 | 7.1 ± 6.8 | 7.1 ± 6.8 | 7.1 ± 6.8 | 7.1 ± 6.8 |
|  | ACER Boltz-2 L-weight | 7.1 ± 6.8 | 7.1 ± 6.8 | 7.1 ± 6.8 | **14.3 ± 9.4** | **14.3 ± 9.4** | **14.3 ± 9.4** |
| 30–40 (23) | Boltz-2 | 26.1 ± 9.1 | 30.4 ± 9.5 | 30.4 ± 9.5 | 30.4 ± 9.4 | 34.8 ± 9.7 | 43.5 ± 10.2 |
|  | Protenix-v1 | **47.8 ± 10.8** | **60.9 ± 10.1** | **69.6 ± 9.4** | **47.8 ± 10.8** | **60.9 ± 10.1** | **69.6 ± 9.4** |
|  | OpenFold-3 | 39.1 ± 10.5 | 47.8 ± 10.4 | 52.2 ± 10.5 | **47.8 ± 10.8** | **60.9 ± 10.2** | 65.2 ± 10.0 |
|  | ACER Boltz-2 L-weight | 17.4 ± 7.9 | 26.1 ± 8.9 | 39.1 ± 10.2 | 30.4 ± 9.5 | 47.8 ± 10.5 | 52.2 ± 10.4 |
| 40–50 (32) | Boltz-2 | **37.5 ± 8.9** | 43.8 ± 9.2 | 46.9 ± 9.2 | **53.1 ± 9.2** | 56.2 ± 9.1 | 59.4 ± 9.1 |
|  | Protenix-v1 | 31.2 ± 8.3 | **53.1 ± 8.9** | **53.1 ± 8.9** | 46.9 ± 8.7 | 62.5 ± 8.6 | 62.5 ± 8.6 |
|  | OpenFold-3 | 34.4 ± 8.9 | 50.0 ± 9.2 | **53.1 ± 9.2** | 46.9 ± 8.8 | 59.4 ± 8.8 | 59.4 ± 8.8 |
|  | ACER Boltz-2 L-weight | **37.5 ± 8.5** | 50.0 ± 9.0 | 50.0 ± 9.0 | **53.1 ± 8.7** | **71.9 ± 8.1** | **71.9 ± 8.1** |
| 50–60 (38) | Boltz-2 | **44.7 ± 8.0** | 47.4 ± 8.1 | 50.0 ± 8.2 | **68.4 ± 7.7** | 68.4 ± 7.7 | **71.1 ± 7.5** |
|  | Protenix-v1 | 36.8 ± 8.0 | 47.4 ± 8.4 | 50.0 ± 8.4 | 36.8 ± 8.0 | 47.4 ± 8.4 | 52.6 ± 8.4 |
|  | OpenFold-3 | 36.8 ± 7.8 | 50.0 ± 7.7 | 55.3 ± 7.8 | 47.4 ± 8.0 | 55.3 ± 7.8 | 57.9 ± 7.7 |
|  | ACER Boltz-2 L-weight | 31.6 ± 7.6 | **55.3 ± 8.1** | **57.9 ± 8.1** | 52.6 ± 8.2 | **71.1 ± 7.5** | **71.1 ± 7.5** |
| 60–70 (38) | Boltz-2 | **57.9 ± 8.0** | 60.5 ± 7.9 | 60.5 ± 7.9 | **63.2 ± 7.7** | 65.8 ± 7.6 | 65.8 ± 7.6 |
|  | Protenix-v1 | 50.0 ± 8.0 | 57.9 ± 8.1 | **63.2 ± 8.0** | 52.6 ± 8.0 | 63.2 ± 7.9 | 65.8 ± 7.9 |
|  | OpenFold-3 | 36.8 ± 7.6 | 42.1 ± 7.8 | 44.7 ± 7.9 | 52.6 ± 8.1 | 57.9 ± 8.1 | 57.9 ± 8.1 |
|  | ACER Boltz-2 L-weight | 55.3 ± 8.2 | **63.2 ± 7.8** | **63.2 ± 7.8** | **63.2 ± 7.8** | **71.1 ± 7.4** | **71.1 ± 7.4** |
| 70–80 (27) | Boltz-2 | **70.4 ± 8.7** | 74.1 ± 8.3 | 74.1 ± 8.3 | **85.2 ± 6.8** | **85.2 ± 6.8** | **85.2 ± 6.8** |
|  | Protenix-v1 | 48.1 ± 9.9 | 74.1 ± 8.7 | 74.1 ± 8.7 | 51.9 ± 9.9 | 77.8 ± 8.3 | 77.8 ± 8.3 |
|  | OpenFold-3 | 40.7 ± 9.4 | 74.1 ± 8.4 | 74.1 ± 8.4 | 44.4 ± 9.5 | 77.8 ± 8.1 | 77.8 ± 8.1 |
|  | ACER Boltz-2 L-weight | 63.0 ± 9.2 | **81.5 ± 7.4** | **81.5 ± 7.4** | 70.4 ± 8.7 | **85.2 ± 6.8** | **85.2 ± 6.8** |
| 80–100 (31) | Boltz-2 | **80.6 ± 6.9** | **80.6 ± 6.9** | **83.9 ± 6.4** | **83.9 ± 6.5** | **83.9 ± 6.5** | **90.3 ± 5.3** |
|  | Protenix-v1 | 48.4 ± 8.7 | 58.1 ± 8.7 | 61.3 ± 8.8 | 54.8 ± 9.0 | 67.7 ± 8.4 | 67.7 ± 8.4 |
|  | OpenFold-3 | 41.9 ± 8.8 | 54.8 ± 8.6 | 58.1 ± 8.6 | 51.6 ± 8.8 | 64.5 ± 8.5 | 67.7 ± 8.4 |
|  | ACER Boltz-2 L-weight | 61.3 ± 8.9 | 74.2 ± 7.9 | 77.4 ± 7.7 | 64.5 ± 8.8 | 80.6 ± 7.2 | 87.1 ± 5.9 |
| **Overall (217)** | Boltz-2 | **47.0 ± 3.4** | 49.8 ± 3.4 | 51.2 ± 3.4 | **57.1 ± 3.4** | 58.5 ± 3.4 | 61.3 ± 3.3 |
|  | Protenix-v1 | 38.2 ± 3.3 | 51.2 ± 3.4 | 53.9 ± 3.4 | 42.4 ± 3.3 | 55.3 ± 3.3 | 57.6 ± 3.3 |
|  | OpenFold-3 | 33.6 ± 3.2 | 46.1 ± 3.3 | 49.3 ± 3.3 | 42.9 ± 3.3 | 54.8 ± 3.3 | 56.2 ± 3.3 |
|  | ACER Boltz-2 L-weight | 40.1 ± 3.4 | **52.5 ± 3.5** | **54.8 ± 3.5** | 50.7 ± 3.5 | **64.1 ± 3.4** | **65.9 ± 3.4** |

### C Equal-compute baselines

ACER runs 30 samples per system, but each carries the overhead of the decoy-augmented trunk and repulsion steering. To match inference cost rather than sample count, we re-run Boltz-2 at the sample count filling the same wall-clock budget: ×150 without steering potentials and ×60 with them, since potentials raise the per-sample cost.

#### C.1 Pocket recovery

**Table C1:** Equal-compute pocket recovery. DCC success rate (%): fraction of systems where at least one predicted pocket falls within 2 Å or 3 Å of the X-ray ligand centre of mass. All rows consume the same inference budget.

| Method | Allosteric ($n = 20$) | | Wrong-pocket ($n = 46$) | |
| --- | --- | --- | --- | --- |
|  | DCC < 2 Å | DCC < 3 Å | DCC < 2 Å | DCC < 3 Å |
| Boltz-2, no potentials ($\times 150$) | 50.0 ± 10.9 | 55.0 ± 10.6 | 34.8 ± 7.1 | 47.8 ± 7.2 |
| Boltz-2, potentials ($\times 60$) | 45.0 ± 11.1 | 50.0 ± 11.2 | 39.1 ± 7.3 | 41.3 ± 7.4 |
| **ACER ($\times 30$)** | **66.6 ± 9.6** | **73.5 ± 9.1** | **44.5 ± 7.1** | **51.7 ± 7.0** |

On pocket recovery benchmark, ACER leads on both benchmarks at both thresholds with the fewest samples. Scaling Boltz-2 to × 150 reaches only 50.0% allosteric recovery at 2 Å, no better than its ×30 configuration (Table 1), and steering potentials do not help. Unsteered samples concentrate in the same dominant pocket however many are drawn, so the gains come from exploration rather than compute.

#### C.2 Pose ranking

ACER’s ensemble-based ranking draws 50 additional poses after pocket exploration, so its total inference cost exceeds that of a single co-folding run at the nominal sample count. To rule out that ACER’s gains come from this extra budget alone, we re-run every baseline with 50 additional samples, matching ACER’s total number of denoiser calls. We report four baseline families: Boltz-2 without steering potentials, Boltz-2 with potentials, Boltz-2 with per-cluster re-ranking (which groups samples by pocket before ranking, isolating the effect of ensemble diversity from ACER’s scoring function), and P2Rank-conditioned Boltz-2. Results are given in Tables C2–C5.

**Table C2:**
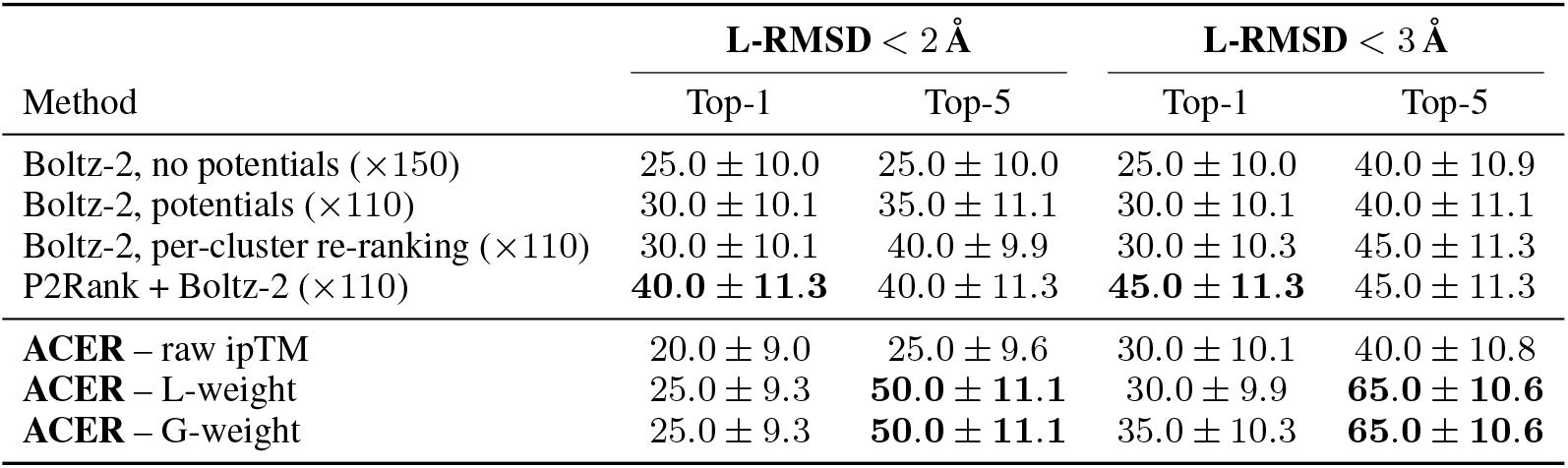
Equal-compute pose ranking on the allosteric set (*n* = 20). Top-*k* L-RMSD success rate (%) at 2 Å and 3 Å, counting only PoseBusters-valid poses (Buttenschoen et al., 2024).

| Method | L-RMSD < 2 Å |  | L-RMSD < 3 Å |  |
| --- | --- | --- | --- | --- |
|  | Top-1 | Top-5 | Top-1 | Top-5 |
| Boltz-2, no potentials ($\times 150$) | 25.0 ± 10.0 | 25.0 ± 10.0 | 25.0 ± 10.0 | 40.0 ± 10.9 |
| Boltz-2, potentials ($\times 110$) | 30.0 ± 10.1 | 35.0 ± 11.1 | 30.0 ± 10.1 | 40.0 ± 11.1 |
| Boltz-2, per-cluster re-ranking ($\times 110$) | 30.0 ± 10.1 | 40.0 ± 9.9 | 30.0 ± 10.3 | 45.0 ± 11.3 |
| P2Rank + Boltz-2 ($\times 110$) | **40.0 ± 11.3** | 40.0 ± 11.3 | **45.0 ± 11.3** | 45.0 ± 11.3 |
| **ACER – raw ipTM** | 20.0 ± 9.0 | 25.0 ± 9.6 | 30.0 ± 10.1 | 40.0 ± 10.8 |
| **ACER – L-weight** | 25.0 ± 9.3 | **50.0 ± 11.1** | 30.0 ± 9.9 | **65.0 ± 10.6** |
| **ACER – G-weight** | 25.0 ± 9.3 | **50.0 ± 11.1** | 35.0 ± 10.3 | **65.0 ± 10.6** |

Under a matched budget, ACER’s advantage is concentrated where pocket identification is the bottleneck, and it is largest at Top-5. On the allosteric set, P2Rank + Boltz-2 attains the best Top-1 (40.0% at 2 Å) but does not improve from Top-1 to Top-5, whereas the ensemble-weighted ACER variants reach 50.0% and 65.0% at Top-5. The same pattern holds on the failed-pocket subset, where ACER leads at both cut-offs. On the full *Runs N’ Poses* set the gap narrows and per-cluster re-ranking is strongest at Top-1, which is expected: most of these systems have a single dominant, well-modelled pocket, so broad exploration has little to recover and ACER’s benefit reduces to its ranking. Restricting to the 0–50 training-similarity bins recovers the advantage, with ACER leading on three of four metrics. Across all four evaluations, increasing the sample count of the baselines does not close the gap on hard or out-of-distribution pockets – additional samples from an unsteered model concentrate in the same dominant site.

**Table C3:** Equal-compute pose ranking on the *Runs N’ Poses* failed-pocket subset (*n* = 46), i.e. systems where Boltz-2 places the ligand in the wrong pocket.

| Method | L-RMSD < 2 Å |  | L-RMSD < 3 Å |  |
| --- | --- | --- | --- | --- |
|  | Top-1 | Top-5 | Top-1 | Top-5 |
| Boltz-2, no potentials ($\times 200$) | 13.0 ± 6.3 | 19.6 ± 6.6 | 19.6 ± 7.3 | 21.7 ± 7.0 |
| Boltz-2, potentials ($\times 110$) | 13.0 ± 6.3 | 17.4 ± 6.7 | 19.6 ± 7.3 | 26.1 ± 7.0 |
| Boltz-2, per-cluster re-ranking ($\times 110$) | 13.0 ± 6.3 | 15.2 ± 6.8 | 19.6 ± 7.1 | 21.7 ± 7.3 |
| P2Rank + Boltz-2 ($\times 110$) | 19.6 ± 5.9 | 26.1 ± 6.6 | 23.9 ± 6.3 | 30.4 ± 6.9 |
| **ACER – raw ipTM** | **23.9 ± 6.4** | 23.9 ± 6.4 | **32.6 ± 7.1** | 34.8 ± 7.2 |
| **ACER – L-weight** | 15.2 ± 5.5 | 26.1 ± 6.7 | 21.7 ± 6.3 | 37.0 ± 7.2 |
| **ACER – G-weight** | 15.2 ± 5.5 | **28.3 ± 6.8** | 21.7 ± 6.3 | **39.1 ± 7.2** |

**Table C4:** Equal-compute pose ranking on the full *Runs N’ Poses* set (*n* = 217).

| Method | L-RMSD < 2 Å |  | L-RMSD < 3 Å |  |
| --- | --- | --- | --- | --- |
|  | Top-1 | Top-5 | Top-1 | Top-5 |
| Boltz-2, no potentials ($\times 200$) | 41.0 ± 3.4 | 49.8 ± 3.4 | 49.8 ± 3.4 | 57.6 ± 3.3 |
| Boltz-2, potentials ($\times 110$) | 47.0 ± 3.4 | 51.6 ± 3.4 | 58.1 ± 3.3 | 61.8 ± 3.2 |
| Boltz-2, per-cluster re-ranking ($\times 110$) | **49.3 ± 3.3** | 49.8 ± 3.3 | **60.8 ± 3.2** | 62.7 ± 3.2 |
| P2Rank + Boltz-2 ($\times 110$) | 37.8 ± 3.3 | 47.5 ± 3.4 | 47.0 ± 3.3 | 56.7 ± 3.3 |
| **ACER – raw ipTM** | 43.8 ± 3.3 | **53.0 ± 3.3** | 54.4 ± 3.4 | 60.4 ± 3.3 |
| **ACER – L-weight** | 40.1 ± 3.4 | 52.5 ± 3.4 | 50.7 ± 3.4 | **64.1 ± 3.3** |

**Table C5:** Equal-compute pose ranking on *Runs N’ Poses* bins. (*n* = 83), comprising systems in the 0–50 training-set similarity bins.

| Method | L-RMSD < 2 Å |  | L-RMSD < 3 Å |  |
| --- | --- | --- | --- | --- |
|  | Top-1 | Top-5 | Top-1 | Top-5 |
| Boltz-2, no potentials ($\times 200$) | 24.1 ± 4.6 | 26.5 ± 4.7 | 26.5 ± 4.7 | 32.5 ± 5.0 |
| Boltz-2, potentials ($\times 110$) | 21.7 ± 4.5 | 25.3 ± 4.7 | 28.9 ± 4.9 | 33.7 ± 5.0 |
| Boltz-2, per-cluster re-ranking ($\times 110$) | **25.3 ± 4.7** | 25.3 ± 4.7 | 33.7 ± 5.3 | 36.1 ± 5.4 |
| P2Rank + Boltz-2 ($\times 110$) | 22.9 ± 4.7 | 28.9 ± 5.1 | 27.7 ± 5.1 | 36.1 ± 5.4 |
| **ACER – raw ipTM** | **25.3 ± 4.6** | **33.7 ± 5.1** | **38.6 ± 5.2** | 42.2 ± 5.3 |
| **ACER – L-weight** | 21.7 ± 4.4 | 28.9 ± 5.1 | 32.5 ± 5.0 | **44.6 ± 5.3** |

##### Runtime, compute and memory overhead

We run ACER primarily on NVIDIA A100-40GB GPUs. For decoy ligand conditioning, the additional cost arises from the extra tokens that must be processed by the trunk module, since triangle attention scales quadratically with token count, inflating both runtime and temporary activation memory. Using 1T49 as a representative structure, Table C6 shows the incremental runtime due to longer trunk processing. Additional diffusion forward passes conditioned on each reconstructed representation compounds runtime overhead accordingly (Table C7).

For iterative pocket repulsion, memory footprint remains unchanged relative to the default, as the input token count is unaffected. However, FK steering incurs extra runtime due to a particle resampling step. Each generated sample under pocket repulsion requires 3× diffusion run time, compared to default sampling (Table C8). Both approaches therefore increase wall-clock time, with the cost scaling jointly with the number of diffusion forward passes and the resampling frequency. As a result, a greater number of exploration iterations leads to proportionally longer runtimes. We also report the overall runtime over Boltz-2 wrong-pocket dataset in Table C9

**Table C6:** Incremental trunk runtime and triangle attention memory footprint as a function of the number of decoy ligands processed by the trunk module, using 1T49 as a representative structure. Trunk cost is estimated as the difference between total forward pass time and diffusion time.

| Number of ligands | Tokens | Tri. Attn., bf16 (GB) | Trunk overhead (s) |
| --- | --- | --- | --- |
| 1 | 286 | 0.17 | $14.0 \pm 0.4$ |
| 5 | 378 | 0.29 | $31.5 \pm 0.1$ |
| 10 | 493 | 0.50 | $71.2 \pm 0.2$ |
| 20 | 608 | 0.76 | $199.3 \pm 0.4$ |

**Table C7:** Runtime comparison between 2 diffusion samples from default sampling and decoy ligand conditioning where each sample was generated in a separate forward pass conditioned on 2 distinct reconstructed representations. Standard deviation is shown over 3 runs.

| Mode | # Samples | Diffusion (s) | Wall-clock runtime (s) |
| --- | --- | --- | --- |
| Boltz-2 default: 1 forward × samples=2 | 2 | $21.2 \pm 0.6$ | $92.3 \pm 8.5$ |
| ACER Boltz-2 – Decoy ligand conditioning: 2 forwards × samples=1 | 2 | $40.8 \pm 0.5$ | $154.0 \pm 1.7$ |

**Table C8:**
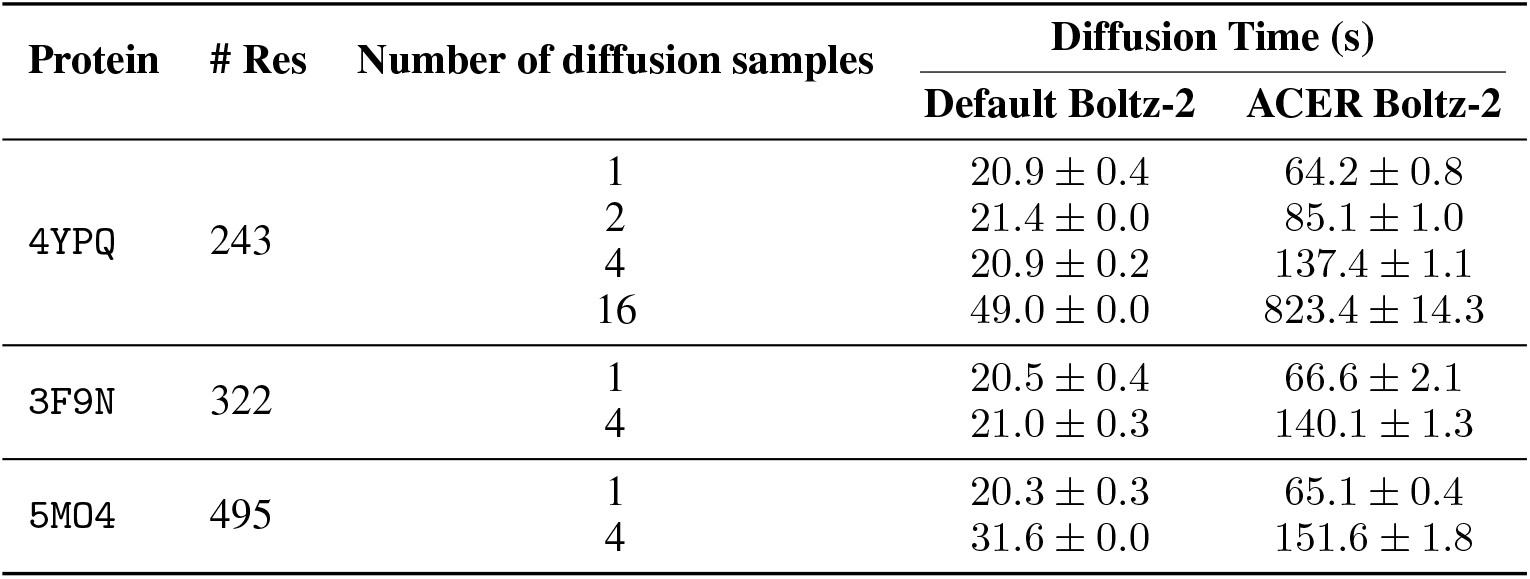
Runtime comparison between Boltz-2 default sampling and ACER with pocket repulsion. Standard deviation is shown over 3 runs.

**Table C9:** Wall-clock runtime breakdown on Boltz-2 wrong-pocket subset for decoy ligand conditioning and iterative pocket repulsion where each diffusion forward pass uses 200 sampling steps. Runtime is reported as mean ± std over benchmark tasks.

| Stage | Config | Sampling steps | Runtime (s) |
| --- | --- | --- | --- |
| Decoy ligand | $n_{\text{ligand}} = 1$ | 200 | $37 \pm 9$ |
| Decoy ligand | $n_{\text{ligand}} = 2$ | 400 | $74 \pm 18$ |
| Decoy ligand | $n_{\text{ligand}} = 3$ | 600 | $111 \pm 27$ |
| Pocket repulsion | $n_{\text{rounds}} = 1$ | 200 | $128 \pm 25$ |
| Pocket repulsion | $n_{\text{rounds}} = 4$ | 800 | $410 \pm 94$ |
| Pocket repulsion | $n_{\text{rounds}} = 5$ | 1000 | $524 \pm 123$ |
| Pocket repulsion | $n_{\text{rounds}} = 8$ | 1600 | $828 \pm 194$ |

### D Ablation study

#### D.1 Pocket exploration

We ablate each component of ACER on the Runs N’ Poses benchmark across 5 seeds (seeds =) to isolate the effect of decoy ligand conditioning and iterative pocket repulsion. All ablations are performed exclusively on the Boltz-2 wrong-pocket subset, allowing us to directly measure the contribution of each ACER component to DCC success rate. Due to the computational costs, we omit the detailed ablation study on ACER – Protenix-v1 expected to yield the similar insights to Boltz-2 case. The results are reported with uncertainty estimates from 95% confidence intervals over 1,000 bootstraps.

As shown in Table D1, decoy ligand conditioning alone yields substantially more correct pocket predictions than iterative pocket repulsion. Notably, pocket repulsion has negligible effect at the strict DCC*<*2Å threshold, suggesting that repulsion guidance in the large search space is insufficient for precise ligand placement. However, by combining both components, ACER attains complementary gains, implying that pocket repulsion becomes effective when the search space is more constrained by occlusion effect introduced by decoy ligand conditioning. For a single seed (seed = 738291) due to computational costs, we therefore ablate the two components with higher iterations of pocket repulsion, demonstrating greater improvements due to increased iterations to substantiate its benefit on top of decoy ligand conditioning (Table D2)

**Table D1:** Ablation of ACER components on the wrong-pocket subset of Runs N’ Poses benchmark for Boltz-2 over 5 seeds (seeds = 231234, 89567, 412903, 451027, 738291)

| Method | Clustered & Distinct Ligands ($n = 46$) | | Total ($n = 88$) | |
| --- | --- | --- | --- | --- |
| | $DCC < 2\text{\AA}$<br>(%)↑ | $DCC < 3\text{\AA}$<br>(%)↑ | $DCC < 2\text{\AA}$<br>(%)↑ | $DCC < 3\text{\AA}$<br>(%)↑ |
| Boltz-2 ($\times 30$ samples) | $30.4 \pm 7.0$ | $37.0 \pm 7.2$ | $36.4 \pm 5.1$ | $43.2 \pm 5.3$ |
| Only decoy ligand conditioning (1–4 decoys) | $43.9 \pm 6.8$ | $50.4 \pm 6.6$ | $42.5 \pm 5.0$ | $48.4 \pm 4.8$ |
| Only 5-round pocket repulsion | $34.8 \pm 6.9$ | $47.8 \pm 7.6$ | $38.6 \pm 4.9$ | $47.7 \pm 5.2$ |
| **Decoy cond. + 5-round pocket repulsion** | **$44.3 \pm 6.5$** | **$51.7 \pm 6.7$** | **$43.6 \pm 5.0$** | **$51.1 \pm 5.1$** |

**Table D2:** Ablation of ACER components on the wrong-pocket subset of Runs N’ Poses benchmark for Boltz-2 on a single seed = 738291.

| Method | Clustered & Distinct Ligands ($n = 46$) | | Total ($n = 88$) | |
| --- | --- | --- | --- | --- |
| | $DCC < 2\text{\AA}$<br>(%)↑ | $DCC < 3\text{\AA}$<br>(%)↑ | $DCC < 2\text{\AA}$<br>(%)↑ | $DCC < 3\text{\AA}$<br>(%)↑ |
| Boltz-2 ($\times 30$ samples) | $30.4 \pm 6.6$ | $37.0 \pm 7.1$ | $36.4 \pm 5.2$ | $43.2 \pm 5.3$ |
| Only decoy ligand conditioning (1–4 decoys) | $45.7 \pm 7.3$ | $50.0 \pm 7.4$ | $43.2 \pm 5.4$ | $52.3 \pm 5.4$ |
| **Decoy cond. + 5-round pocket repulsion** | **$45.7 \pm 7.2$** | **$52.2 \pm 7.4$** | **$43.2 \pm 5.2$** | **$54.5 \pm 5.4$** |
| **Decoy cond. + 15-round pocket repulsion** | **$47.8 \pm 7.6$** | **$54.3 \pm 7.2$** | **$46.6 \pm 5.2$** | **$58.0 \pm 5.1$** |

Table D3 shows the effect of the number of decoy ligands on the conditioning representation. Performance improves consistently with more decoys across both thresholds and both subsets, suggesting that a larger ensemble of decoy poses occludes more of the surface around the incorrect pocket, steering the model toward alternative binding interfaces that are underexplored in default sampling mode.

**Table D3:** Ablation as a function of the number of decoy ligands on the Runs N’ Poses and Allosteric Systems benchmarks on a single seed = 738291.

| # Total ligands | # Decoy ligands | Clustered & Distinct Ligands ($n = 46$) | | Total ($n = 88$) | | Allosteric Systems ($n = 20$) | |
| --- | --- | --- | --- | --- | --- | --- | --- |
| | | $DCC < 2\text{\AA}$<br>(%)↑ | $DCC < 3\text{\AA}$<br>(%)↑ | $DCC < 2\text{\AA}$<br>(%)↑ | $DCC < 3\text{\AA}$<br>(%)↑ | $DCC < 2\text{\AA}$<br>(%)↑ | $DCC < 3\text{\AA}$<br>(%)↑ |
| 2 | 1 | $41.4 \pm 7.3$ | $47.8 \pm 7.4$ | $36.9 \pm 5.1$ | $46.6 \pm 5.3$ | $44.7 \pm 11.2$ | $50.0 \pm 10.7$ |
| 3 | 2 | $45.6 \pm 7.4$ | $50.5 \pm 7.2$ | $41.2 \pm 5.3$ | $48.8 \pm 5.4$ | $54.4 \pm 10.8$ | $59.6 \pm 10.9$ |
| 4 | 3 | **$45.9 \pm 7.5$** | $49.9 \pm 7.3$ | $42.3 \pm 5.1$ | $50.2 \pm 5.3$ | $55.1 \pm 11.5$ | $60.2 \pm 10.9$ |
| 5 | 4 | $45.8 \pm 7.5$ | **$50.3 \pm 7.3$** | **$43.3 \pm 5.5$** | **$51.3 \pm 5.2$** | **$55.1 \pm 11.2$** | **$65.4 \pm 10.6$** |

Table D4 shows DCC success rates from decoy ligand conditioning, accumulated across decoy ligand types for 1-4 decoy ligands and 1 target ligand. Using decoy ligands identical to the target ligand performs comparably to small and medium chemical probes (KI2 and FER). In contrast, the bulkier PTY probe underperforms, likely because its steric bulk constrains the range of plausible binding sites. Since chemical probes offer no clear advantage in exploration, we use native ligands as decoy ligands in this work, as their duplicate representations for conditioning can be cached once and reused across multiple diffusion runs.

**Table D4:** Ablation of the choice of decoy ligand type on DCC success rate (%) at 2 Å and 3 Å thresholds.

| Decoy ligand type | Clustered & Distinct Ligands ($n = 46$) | | Total ($n = 88$) | |
| --- | --- | --- | --- | --- |
|  | DCC<2 Å<br>(%)↑ | DCC<3 Å<br>(%)↑ | DCC<2 Å<br>(%)↑ | DCC<3 Å<br>(%)↑ |
| Native (5× native) | **45.3 ± 7.2** | 50.2 ± 7.3 | **43.2 ± 5.1** | **52.6 ± 5.2** |
| KI2 | 26.1 ± 6.6 | **35.1 ± 6.8** | 29.7 ± 5.0 | 38.5 ± 5.4 |
| PTY | 15.2 ± 5.3 | 28.3 ± 6.7 | 21.7 ± 4.5 | 34.0 ± 5.1 |
| FER | 25.9 ± 6.5 | 30.3 ± 6.6 | 25.1 ± 4.6 | 31.8 ± 4.9 |

We ablate the pocket repulsion algorithm parameters without decoy ligand conditioning to assess how DCC success rate is sensitive to the parameters in guidance sampling, including the virtual pocket radius *r*_*g*_ (Table D5), the minimum distance threshold *d*_min_ (Table D6), and optional protein-ligand contact guidance that preserves protein-ligand contact (Table D7). The results are derived from a single seed (seed = 738291). Uncertainty is estimated by 1000 bootstrap resampling.

Table D5 reports DCC success rate by varying the virtual pocket radius *r*_*g*_ used to define the repulsion region. Larger *r*_*g*_ improve the likelihood of correct ligand placement into alternative pockets, with *r* = 8 achieving the best results. This suggests that larger repulsion against the already explored regions is more effective to discover the new binding interface.

**Table D5:** Ablation as a function of the pocket radius on the Runs N’ Poses benchmark over 5 iterations of pocket repulsion. Boltz-2 baseline runs default sampling without decoy ligand conditioning

| Pocket radius | Clustered & Distinct Ligands ($n = 46$) | | Total ($n = 88$) | |
| --- | --- | --- | --- | --- |
|  | DCC<2 Å<br>(%)↑ | DCC<3 Å<br>(%)↑ | DCC<2 Å<br>(%)↑ | DCC<3 Å<br>(%)↑ |
| Boltz-2 (×5 samples without pocket repulsion) | 26.1 ± 6.5 | 34.9 ± 7.4 | 30.6 ± 5.1 | 36.6 ± 5.2 |
| 4 Å | 28.4 ± 6.5 | 32.4 ± 6.8 | 34.1 ± 5.2 | 38.6 ± 4.9 |
| 6 Å | **32.5 ± 6.8** | 36.8 ± 7.1 | **36.4 ± 4.8** | 40.9 ± 5.2 |
| 8 Å | 28.3 ± 6.8 | **41.4 ± 7.4** | 31.7 ± 5.1 | **40.9 ± 5.3** |

Table D6 shows DCC success rate by the minimum distance threshold *d*_min_ controlling how far decoy ligands must be placed from the restricting binidng pocket. Performance degrades at smaller values (*d*_min_ ∈ {4, 6}), indicating that ligands placed too close to the previous pocket do not effectively explore alternative pockets.

As applying the repulsion potential alone risks pushing the ligand entirely away from the protein surface, we experiment introducing the contact potential *U*_protein-contact_ in Equation D12 as a safeguard to keep the ligand in contact with protein but not so close to cause steric clashes. We therefore ablate the contact weight added to the existing potential where *λ*_c_ to quantify its effect. Table D7 reports the effect of varying the contact weight hyperparameter. Disabling contact weighting entirely (*λ*_c_ = 0.0) performs similarly on DCC success rate *<* 2 Å threshold, suggesting that conditioning representations that direct the denoiser already implicitly enforce structural priors to maintain protein-ligand contact.

The protein-ligand contact potential provides limited benefit in this setting.

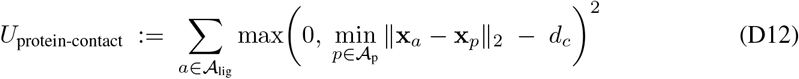

where *A*_lig_ and *A*_p_ denote the sets of ligand and protein heavy atoms, respectively, **x**_*p*_ ∈ ℝ^3^ is the position of protein atom *p*, and *d*_*c*_ is the maximum allowed distance to the nearest protein atom. The term on added to the existing potentials ^3^.

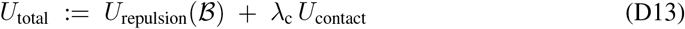

where *λ*_c_ ≥ 0 is a scalar weight controlling the strength of the contact term. Setting *λ*_c_ = 0 refers to pure repulsion steering.

**Table D6:** Ablation as a function of the minimum distance threshold *d*_min_ on the Runs N’ Poses benchmark over 5 iterations of pocket repulsion. Boltz-2 baseline runs default sampling without decoy ligand conditioning

| $d_{\min}$ | Clustered & Distinct Ligands ($n = 46$) | | Total ($n = 88$) | |
| --- | --- | --- | --- | --- |
|  | DCC<2 Å<br>(%)↑ | DCC<3 Å<br>(%)↑ | DCC<2 Å<br>(%)↑ | DCC<3 Å<br>(%)↑ |
| Boltz-2 ($\times 5$ samples without pocket repulsion) | 26.1 ± 6.5 | 34.9 ± 7.4 | 30.6 ± 5.1 | 36.6 ± 5.2 |
| 4 Å | 23.8 ± 6.2 | 32.4 ± 7.0 | 30.7 ± 4.8 | 35.5 ± 5.0 |
| 6 Å | 28.1 ± 6.9 | 34.9 ± 7.0 | **34.9 ± 5.3** | 38.4 ± 4.9 |
| 8 Å | **28.1 ± 6.8** | **41.6 ± 7.5** | 31.7 ± 4.9 | **41.0 ± 5.3** |

**Table D7:** Ablation as a function of the contact weight on the Runs N’ Poses benchmark over 5 iterations of pocket repulsion. Boltz-2 baseline runs default sampling without decoy ligand conditioning.

| Contact weight | Clustered & Distinct Ligands ($n = 46$) | | Total ($n = 88$) | |
| --- | --- | --- | --- | --- |
|  | DCC<2 Å<br>(%)↑ | DCC<3 Å<br>(%)↑ | DCC<2 Å<br>(%)↑ | DCC<3 Å<br>(%)↑ |
| Boltz-2 ($\times 5$ samples without pocket repulsion) | 26.1 ± 6.5 | 34.9 ± 7.4 | 30.6 ± 5.1 | 36.6 ± 5.2 |
| 0.0 | **30.7 ± 7.0** | 41.5 ± 7.1 | **34.3 ± 5.3** | 40.8 ± 5.3 |
| 0.5 | 28.2 ± 6.6 | 38.8 ± 7.3 | 33.1 ± 5.0 | **40.9 ± 5.5** |
| 1.0 | 27.8 ± 6.9 | **41.7 ± 7.4** | 31.9 ± 5.0 | 40.6 ± 5.3 |

#### D.2 Ensemble-based ranking

We ablate how the two key sampling-budget parameters affect ensemble-based pose ranking across the allosteric set, the Boltz-2 wrong-pocket subset of *Runs N’ Poses*, and the full clustered-and-distinct ligand subset. Uncertainty is reported as 95% confidence intervals over 1,000 bootstraps.

Given the total sampling budget *B* = *P* × *S, P* is the number of candidate binding pockets obtained from the exploration phase, while *S* represents the local conformational coverage within each pocket. We ablate both parameters across all test sets in Tables D8–D11. A modest budget of *B* = 50 samples is typically sufficient for ACER to outperform the baselines. However, on the generalizability benchmark, increasing the budget to *B* = 200 yields substantially higher success rates in the hardest similarity bin (≤ 20%), where pocket discovery is the key contributor and retaining more candidate pockets in the ranking pool becomes critical (Figure 2).

**Table D8:** Ablation on selected *P* pockets and *S* local ensemble conformers within the pocket on allosteric set (*n* = 20).

| Ranking Approach | $P$ | $S$ | % L-RMSD < 2 Å & PB Valid ↑ | | | % L-RMSD < 3 Å & PB Valid ↑ | | |
| --- | --- | --- | --- | --- | --- | --- | --- | --- |
|  |  |  | Top-1 | Top-5 | Top-10 | Top-1 | Top-5 | Top-10 |
| *Budget = 50* |  |  |  |  |  |  |  |  |
| Boltz-2 | 10 | 5 | 25.0 ± 9.6 | 30.0 ± 10.1 | 30.0 ± 10.1 | 30.0 ± 10.2 | 40.0 ± 11.0 | 40.0 ± 11.0 |
| ACER – baseline | 10 | 5 | 20.0 ± 9.0 | 25.0 ± 9.6 | 35.0 ± 10.6 | 30.0 ± 10.1 | 40.0 ± 10.8 | 45.0 ± 10.9 |
| ACER – L-weight | 10 | 5 | **25.0 ± 9.3** | **50.0 ± 11.1** | **50.0 ± 11.1** | 30.0 ± 9.9 | **65.0 ± 10.6** | **70.0 ± 10.3** |
| ACER – G-weight | 10 | 5 | **25.0 ± 9.3** | **50.0 ± 11.1** | **50.0 ± 11.1** | **35.0 ± 10.3** | **65.0 ± 10.6** | **70.0 ± 10.3** |
| *Budget = 100* |  |  |  |  |  |  |  |  |
| Boltz-2 | 10 | 10 | 25.0 ± 9.6 | 30.0 ± 10.1 | 30.0 ± 10.1 | 30.0 ± 10.2 | 40.0 ± 11.0 | 40.0 ± 11.0 |
| ACER – L-weight | 10 | 10 | **30.0 ± 10.1** | **50.0 ± 11.2** | **55.0 ± 11.1** | **35.0 ± 10.6** | 60.0 ± 11.0 | **70.0 ± 10.3** |
| ACER – G-weight | 10 | 10 | **30.0 ± 10.1** | **50.0 ± 11.2** | **55.0 ± 11.1** | **35.0 ± 10.6** | **65.0 ± 11.0** | **70.0 ± 10.3** |
| *Budget = 200* |  |  |  |  |  |  |  |  |
| Boltz-2 | 20 | 10 | 25.0 ± 9.6 | 30.0 ± 10.1 | 30.0 ± 10.1 | 30.0 ± 10.2 | 40.0 ± 11.0 | 40.0 ± 11.0 |
| ACER – L-weight | 20 | 10 | 25.0 ± 9.6 | **45.0 ± 11.4** | **55.0 ± 11.1** | 30.0 ± 10.2 | 60.0 ± 11.0 | **70.0 ± 10.3** |
| ACER – G-weight | 20 | 10 | 25.0 ± 9.6 | **45.0 ± 11.4** | **55.0 ± 11.1** | 30.0 ± 10.2 | **65.0 ± 11.0** | **70.0 ± 10.3** |

**Table D9:** Ablation on selected *P* pockets and *S* local ensemble conformers within the pocket on *Runs N’ Poses* wrong-pocket subset of Boltz-2 – Clustered & Distinct Ligands (*n* = 46).

| Ranking Approach | $P$ | $S$ | % L-RMSD < 2 Å & PB Valid ↑ | | % L-RMSD < 3 Å & PB Valid ↑ | |
| --- | --- | --- | --- | --- | --- | --- |
|  |  |  | Top-1 | Top-5 | Top-1 | Top-5 |
| *Budget = 50* |  |  |  |  |  |  |
| Boltz-2 | 10 | 5 | 15.2 ± 5.5 | 15.2 ± 5.5 | 19.6 ± 6.0 | 19.6 ± 6.0 |
| ACER – baseline | 10 | 5 | **23.9 ± 6.4** | 23.9 ± 6.4 | **32.6 ± 7.1** | 34.8 ± 7.2 |
| ACER – L-weight | 10 | 5 | 15.2 ± 5.5 | 26.1 ± 6.7 | 21.7 ± 6.3 | 37.0 ± 7.2 |
| ACER – G-weight | 10 | 5 | 15.2 ± 5.5 | **28.3 ± 6.8** | 21.7 ± 6.3 | **39.1 ± 7.2** |
| *Budget = 100* |  |  |  |  |  |  |
| Boltz-2 | 10 | 10 | 15.2 ± 5.5 | 15.2 ± 5.5 | 19.6 ± 6.0 | 19.6 ± 6.0 |
| ACER – baseline | 10 | 10 | **23.9 ± 6.4** | 23.9 ± 6.4 | **32.6 ± 7.1** | 34.8 ± 7.2 |
| ACER – L-weight | 10 | 10 | 19.6 ± 6.0 | **28.3 ± 7.0** | 28.3 ± 6.8 | **39.1 ± 7.4** |
| ACER – G-weight | 10 | 10 | 19.6 ± 6.0 | 26.1 ± 6.7 | 28.3 ± 6.8 | 37.0 ± 7.4 |
| *Budget = 200* |  |  |  |  |  |  |
| Boltz-2 | 20 | 10 | 15.2 ± 5.5 | 15.2 ± 5.5 | 19.6 ± 6.0 | 19.6 ± 6.0 |
| ACER – baseline | 20 | 10 | 21.7 ± 6.2 | 23.9 ± 6.4 | 28.3 ± 6.7 | 34.8 ± 7.2 |
| ACER – L-weight | 20 | 10 | 21.7 ± 6.2 | **26.1 ± 6.8** | **32.6 ± 7.1** | **39.1 ± 7.4** |
| ACER – G-weight | 20 | 10 | 21.7 ± 6.2 | 23.9 ± 6.5 | **32.6 ± 7.1** | 37.0 ± 7.4 |

**Table D10:** Ablation on selected *P* pockets and *S* local ensemble conformers within the pocket on the *Runs N’ Poses* wrong-pocket subset of Boltz-2 – Total (*n* = 88).

| Method | $P$ | $S$ | % L-RMSD < 2 Å & PB Valid ↑ | | % L-RMSD < 3 Å & PB Valid ↑ | |
| --- | --- | --- | --- | --- | --- | --- |
|  |  |  | Top-1 | Top-5 | Top-1 | Top-5 |
| *Budget = 50* |  |  |  |  |  |  |
| Boltz-2 | 10 | 5 | 26.1 ± 4.8 | 27.3 ± 4.8 | 28.4 ± 4.9 | 29.5 ± 5.0 |
| ACER – baseline | 10 | 5 | **30.7 ± 5.0** | 31.8 ± 5.1 | **35.2 ± 5.2** | 37.5 ± 5.3 |
| ACER – L-weight | 10 | 5 | 21.6 ± 4.4 | 29.5 ± 4.9 | 26.1 ± 4.8 | 36.4 ± 5.3 |
| ACER – G-weight | 10 | 5 | 21.6 ± 4.4 | **31.8 ± 5.1** | 26.1 ± 4.8 | **38.6 ± 5.4** |
| *Budget = 100* |  |  |  |  |  |  |
| Boltz-2 | 10 | 10 | 26.1 ± 4.8 | 27.3 ± 4.8 | 28.4 ± 4.9 | 29.5 ± 5.0 |
| ACER – baseline | 10 | 10 | **30.7 ± 5.0** | 31.8 ± 5.1 | **35.2 ± 5.2** | 37.5 ± 5.3 |
| ACER – L-weight | 10 | 10 | 25.0 ± 4.7 | 30.7 ± 5.0 | 29.5 ± 5.0 | **37.5 ± 5.3** |
| ACER – G-weight | 10 | 10 | 25.0 ± 4.7 | 29.5 ± 4.9 | 29.5 ± 5.0 | 36.4 ± 5.3 |
| *Budget = 200* |  |  |  |  |  |  |
| Boltz-2 | 20 | 10 | 26.1 ± 4.8 | 27.3 ± 4.8 | 28.4 ± 4.9 | 29.5 ± 5.0 |
| ACER – baseline | 20 | 10 | **29.5 ± 5.0** | **33.0 ± 5.2** | **33.0 ± 5.2** | **38.6 ± 5.4** |
| ACER – L-weight | 20 | 10 | 26.1 ± 4.7 | 29.5 ± 4.9 | 31.8 ± 5.1 | 37.5 ± 5.3 |
| ACER – G-weight | 20 | 10 | 26.1 ± 4.7 | 28.4 ± 4.8 | 31.8 ± 5.1 | 36.4 ± 5.3 |

**Table D11:**
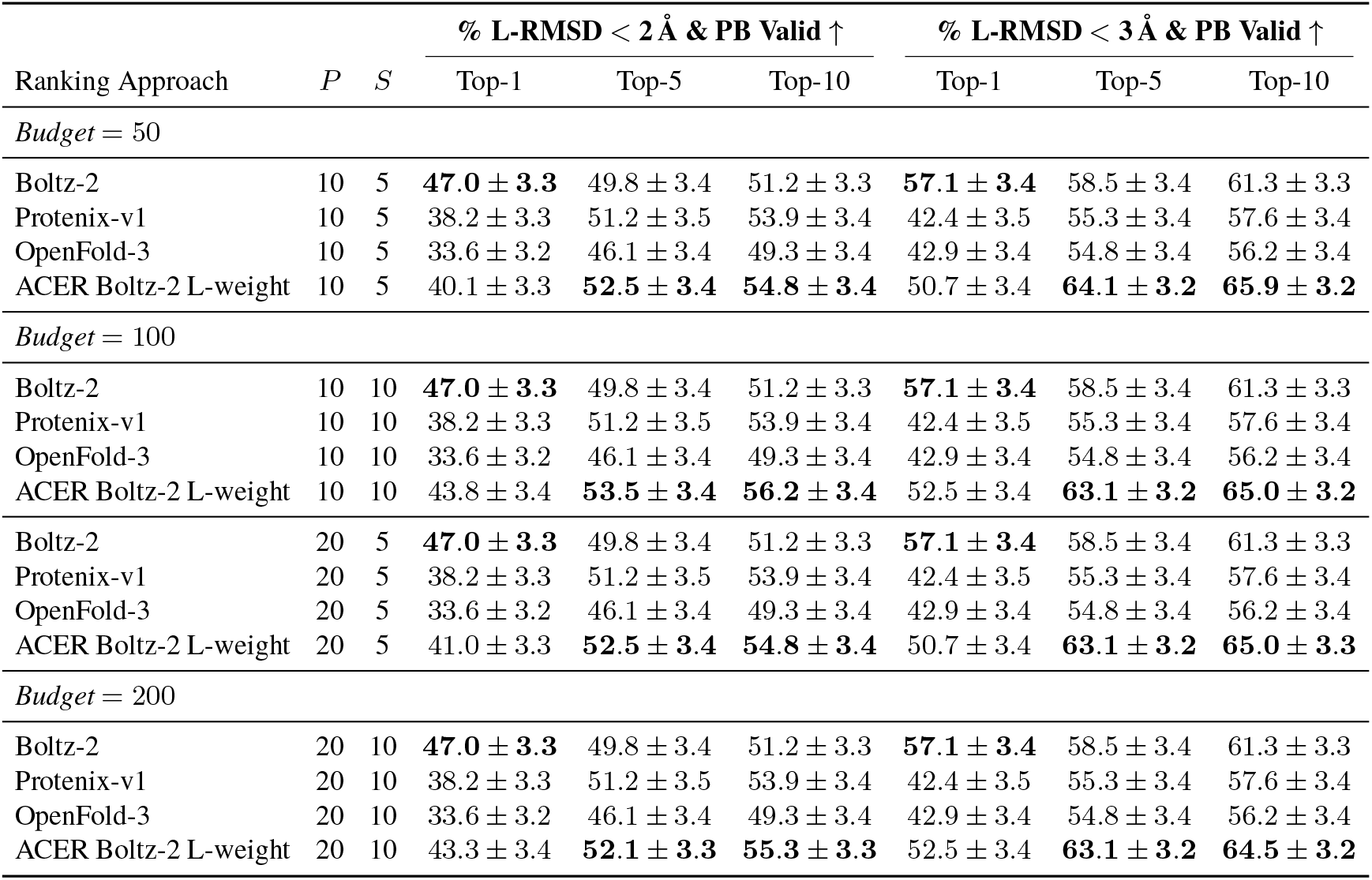
Ablation on selected *P* pockets and *S* local ensemble conformers within the pocket on the full scope of *Runs N’ Poses* Clustered and Distinct Ligands (*n* = 217).

Algorithm 1 details the iterative pocket repulsion procedure. Over *R* sampling rounds, the model accumulates a set of occupied pockets *B*, from a prior set of pockets to steer away from (e.g., known prevalent or previously sampled pockets). In each round, a pose is sampled under the repulsion potential *U*_repulsion_(*ℬ*), which penalises all pockets in *ℬ* and steers the model toward unexplored regions. The pocket residues *ℛ*_*r*_ for the resulting pose are extracted by selecting all residues with any heavy atom within 6 Å of any ligand atom, and added to *ℬ* to repel subsequent rounds. User-specified parameters include the number of rounds *R*, the distance threshold for pocket extraction, and the choice of prior pockets in *ℬ*_0_.

##### Algorithm 1

Iterative pocket repulsion

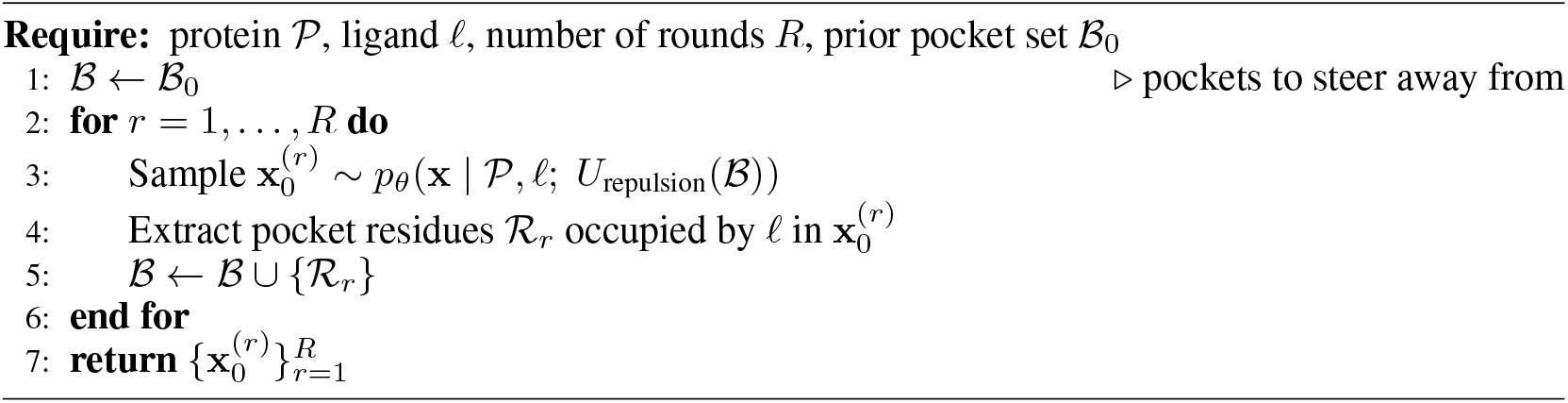

**Figure D1:**
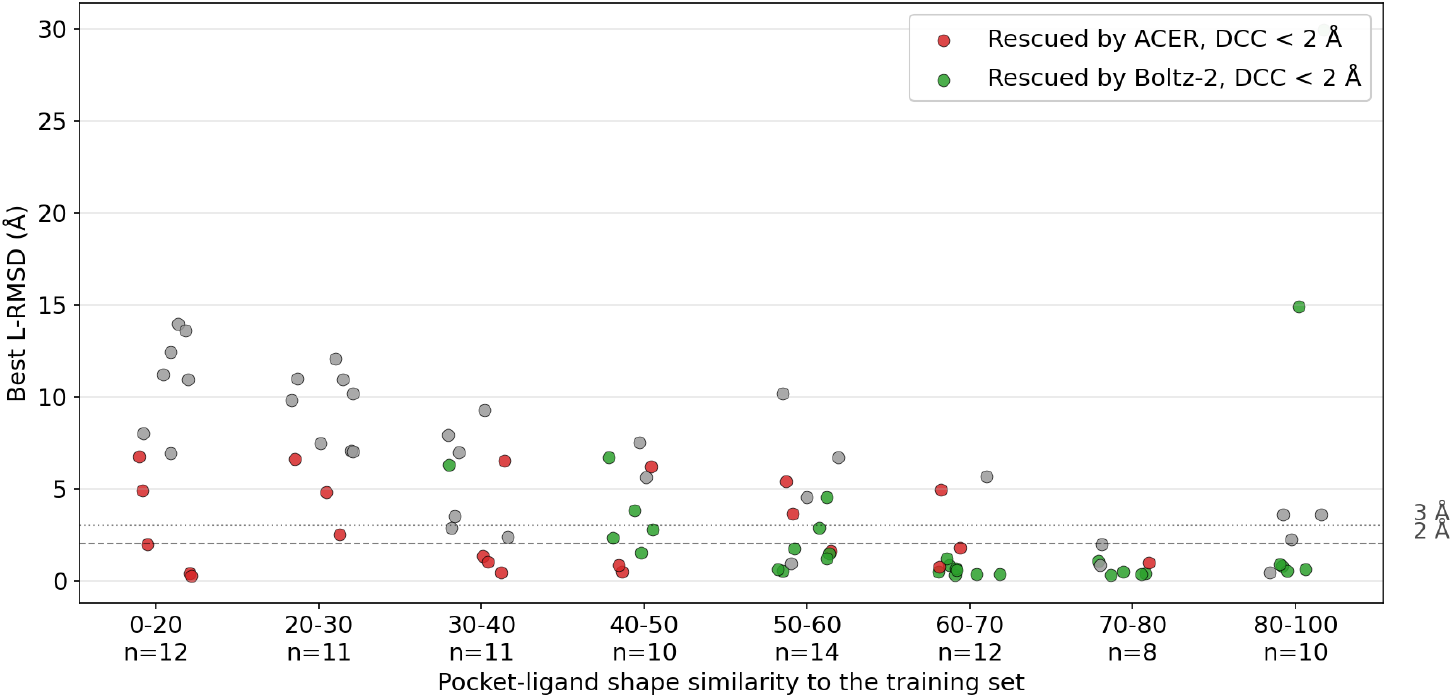
Best L-RMSD per system, stratified by pocket-ligand shape similarity to the training set. Red indicates systems where the Boltz-2 baseline fails (DCC ≥ 2 Å) but ACER rescues the correct pocket (DCC *<* 2 Å); green indicates systems where Boltz-2 alone already succeeds. Gray dots indicate systems where both methods fail. ACER rescue is most prevalent at low similarity (0–30), where Boltz-2 consistently fails to place ligands, and can sample the accurate poses (L-RMSD below 2 Å) in several cases.

**Figure D2:**
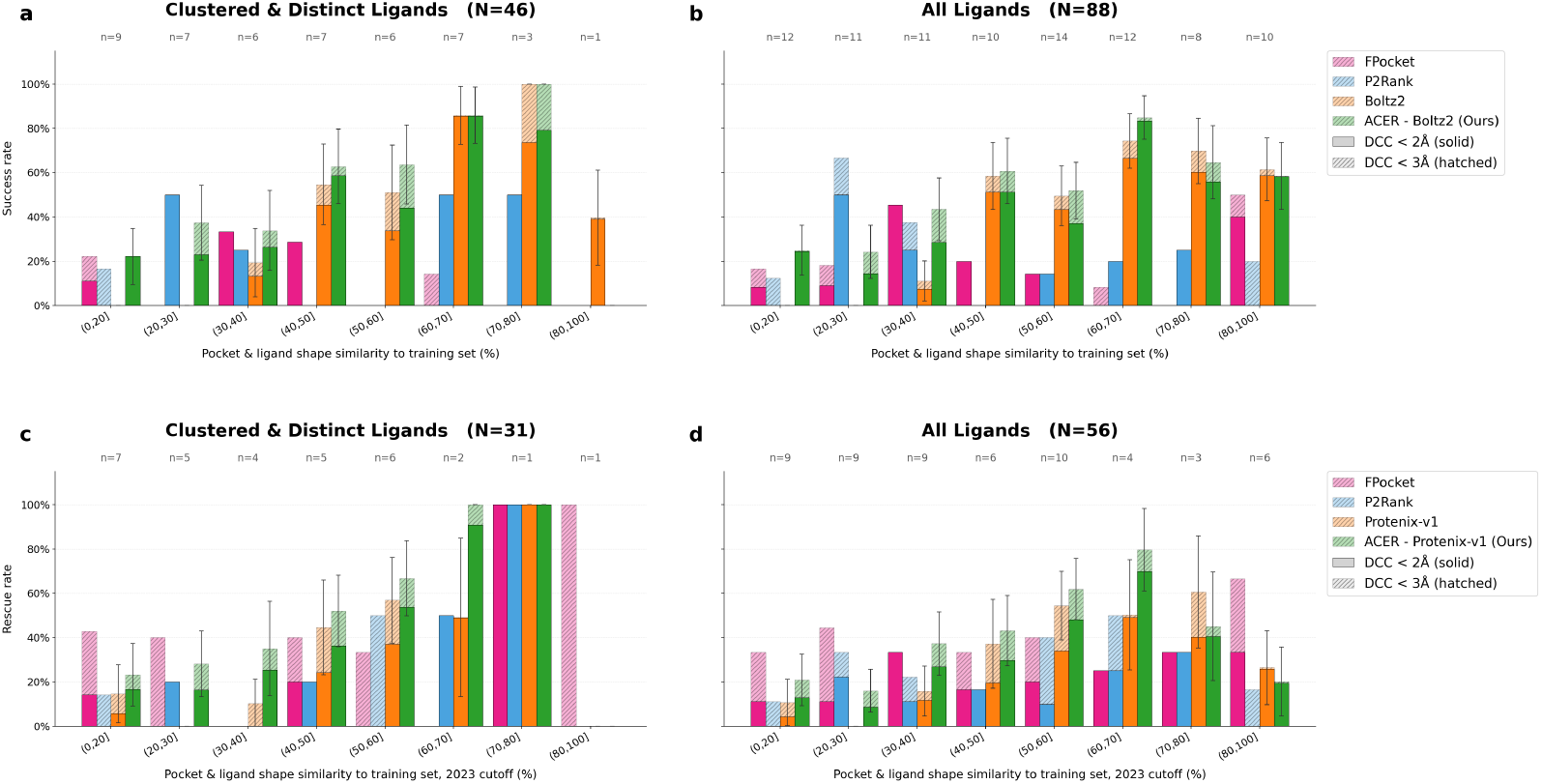
DCC success rate on the wrong-pocket subset of Runs N’ Poses (after 2023-06-01), stratified by pocket–ligand shape similarity to the training set. **(a, b)** Systems where top-ranked Boltz-2 misplaces the ligand (DCC *>* 2 Å). **(c, d)** Same, but for Protenix-v1. Filtered (*n* = 46, *n* = 31) and full (*n* = 88, *n* = 55) subsets shown in the left and right columns, respectively.

**Figure D3:**
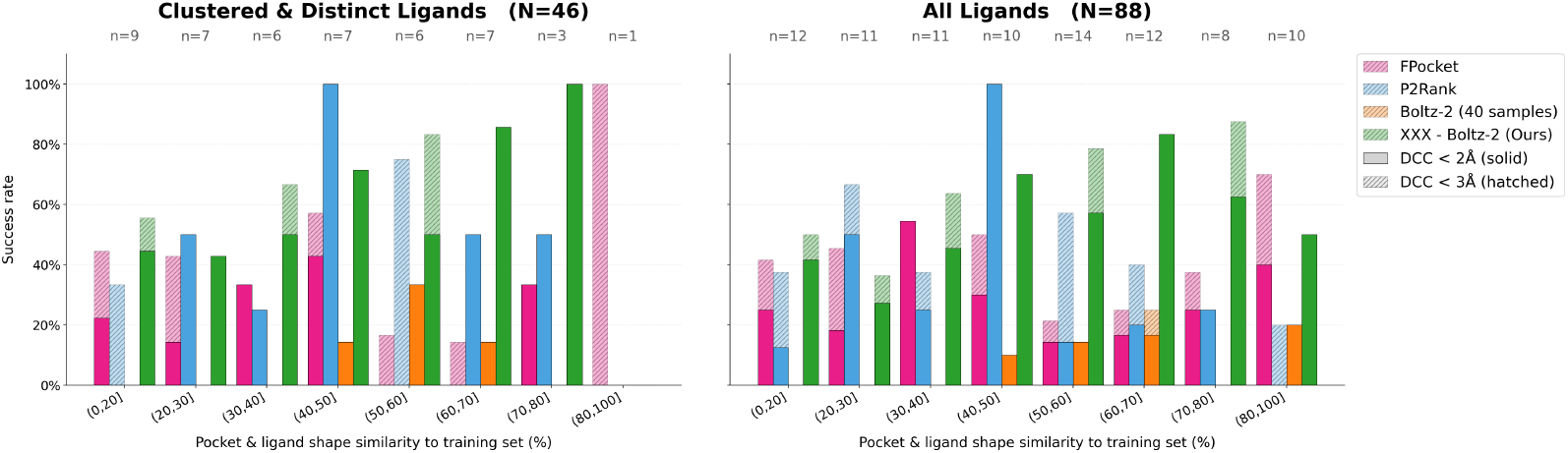
DCC success rate on the wrong-pocket subset of Runs N’ Poses when considering oracle across 5 seeds. The system is considered successful if at least one prediction across 5 seeds achieves a DCC below the.

**Figure D4:**
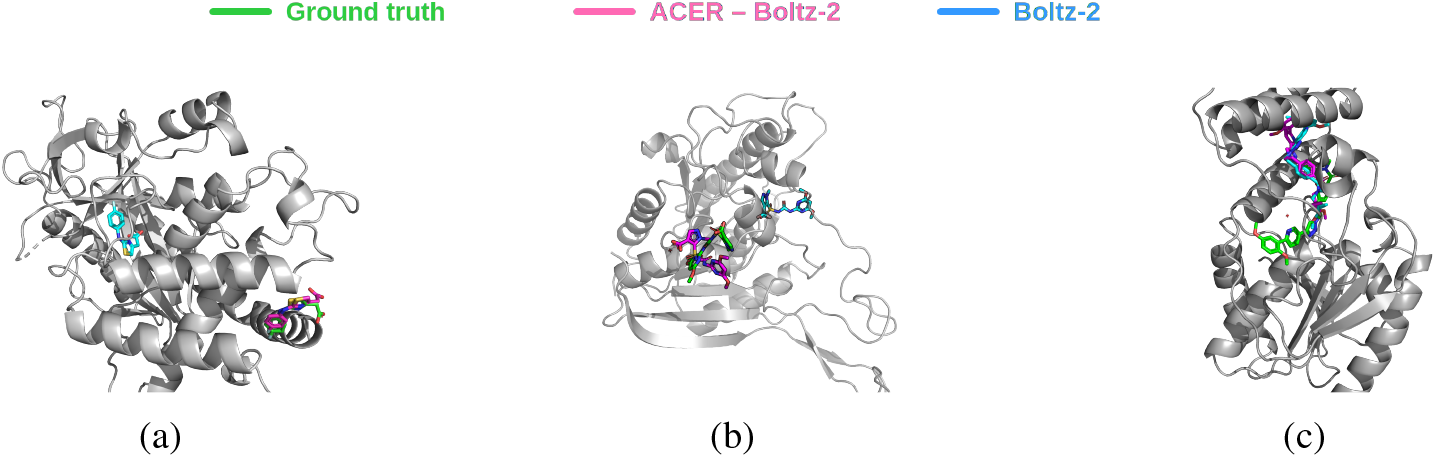
Qualitative examples in the (0, 20] similarity bin of the Runs N’ Poses benchmark. (a) 7FTM – where ACER correctly places ligands into the right pocket (DCC *<* 2 Å) *and* recovers an accurate ligand pose (L-RMSD *<* 2 Å). (b) 8GOY – a partial success where ligand placement succeeds (DCC *<* 2 Å), yet the predicted ligand pose remains inaccurate (L-RMSD *>* 2 Å). (c) 8TQV – a failure case in which both ACER – Boltz-2 and default Boltz-2 can never rescue the correct pocket.

**Figure D5:**
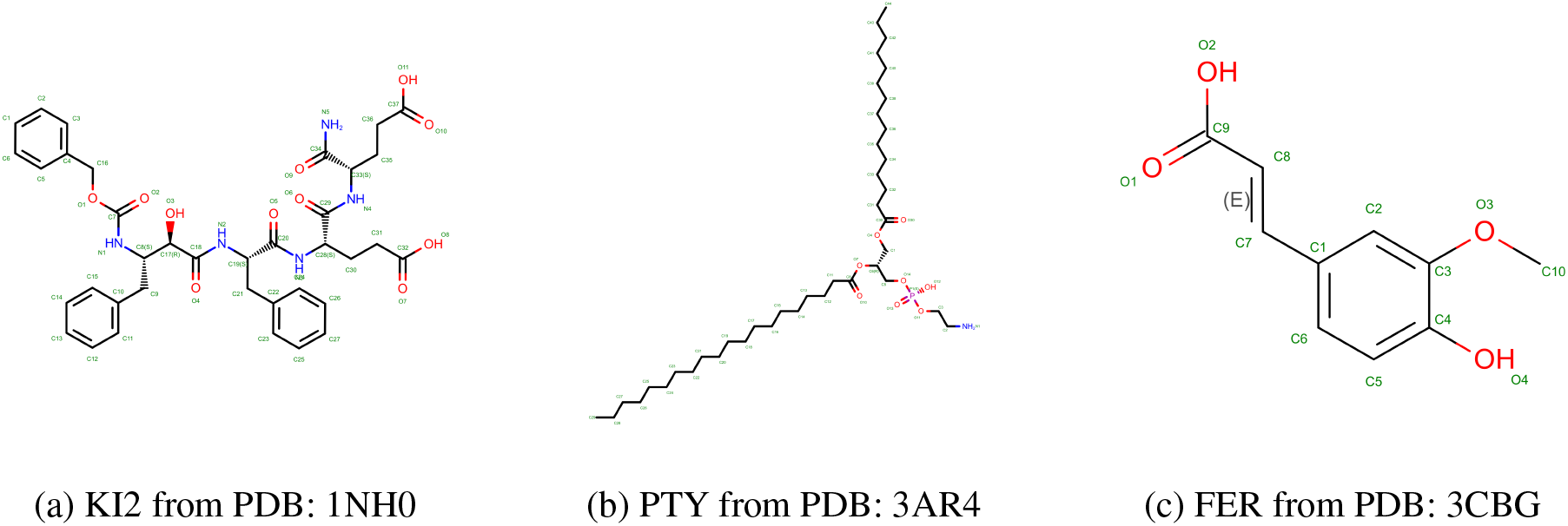
The structure of chemical probes used in the ablation study of decoy ligand type.

##### NeurIPS Paper Checklist

1. **Claims** Question: Do the main claims made in the abstract and introduction accurately reflect the paper’s contributions and scope? Answer: [Yes] Justification: We believe that the claims of the abstract and introduction are fully supported by the paper text. Guidelines:
  - The answer [N/A] means that the abstract and introduction do not include the claims made in the paper.
  - The abstract and/or introduction should clearly state the claims made, including the contributions made in the paper and important assumptions and limitations. A [No] or [N/A] answer to this question will not be perceived well by the reviewers.
  - The claims made should match theoretical and experimental results, and reflect how much the results can be expected to generalize to other settings.
  - It is fine to include aspirational goals as motivation as long as it is clear that these goals are not attained by the paper.
2. **Limitations** Question: Does the paper discuss the limitations of the work performed by the authors? Answer: [Yes] Justification: See Limitations section. Guidelines:
  - The answer [N/A] means that the paper has no limitation while the answer [No] means that the paper has limitations, but those are not discussed in the paper.
  - The authors are encouraged to create a separate “Limitations” section in their paper.
  - The paper should point out any strong assumptions and how robust the results are to violations of these assumptions (e.g., independence assumptions, noiseless settings, model well-specification, asymptotic approximations only holding locally). The authors should reflect on how these assumptions might be violated in practice and what the implications would be.
  - The authors should reflect on the scope of the claims made, e.g., if the approach was only tested on a few datasets or with a few runs. In general, empirical results often depend on implicit assumptions, which should be articulated.
  - The authors should reflect on the factors that influence the performance of the approach. For example, a facial recognition algorithm may perform poorly when image resolution is low or images are taken in low lighting. Or a speech-to-text system might not be used reliably to provide closed captions for online lectures because it fails to handle technical jargon.
  - The authors should discuss the computational efficiency of the proposed algorithms and how they scale with dataset size.
  - If applicable, the authors should discuss possible limitations of their approach to address problems of privacy and fairness.
  - While the authors might fear that complete honesty about limitations might be used by reviewers as grounds for rejection, a worse outcome might be that reviewers discover limitations that aren’t acknowledged in the paper. The authors should use their best judgment and recognize that individual actions in favor of transparency play an important role in developing norms that preserve the integrity of the community. Reviewers will be specifically instructed to not penalize honesty concerning limitations.
3. **Theory assumptions and proofs** Question: For each theoretical result, does the paper provide the full set of assumptions and a complete (and correct) proof? Answer: [N/A] Justification: The paper does not include theoretical results. However, the associated theories are cited. Guidelines:
  - The answer [N/A] means that the paper does not include theoretical results.
  - All the theorems, formulas, and proofs in the paper should be numbered and crossreferenced.
  - All assumptions should be clearly stated or referenced in the statement of any theorems.
  - The proofs can either appear in the main paper or the supplemental material, but if they appear in the supplemental material, the authors are encouraged to provide a short proof sketch to provide intuition.
  - Inversely, any informal proof provided in the core of the paper should be complemented by formal proofs provided in appendix or supplemental material.
  - Theorems and Lemmas that the proof relies upon should be properly referenced.
4. **Experimental result reproducibility** Question: Does the paper fully disclose all the information needed to reproduce the main experimental results of the paper to the extent that it affects the main claims and/or conclusions of the paper (regardless of whether the code and data are provided or not)? Answer: [Yes] Justification: Details of experimental design and algorithms are discussed in details in the paper and Supplementary Materials. Guidelines:
  - The answer [N/A] means that the paper does not include experiments.
  - If the paper includes experiments, a [No] answer to this question will not be perceived well by the reviewers: Making the paper reproducible is important, regardless of whether the code and data are provided or not.
  - If the contribution is a dataset and/or model, the authors should describe the steps taken to make their results reproducible or verifiable.
  - Depending on the contribution, reproducibility can be accomplished in various ways. For example, if the contribution is a novel architecture, describing the architecture fully might suffice, or if the contribution is a specific model and empirical evaluation, it may be necessary to either make it possible for others to replicate the model with the same dataset, or provide access to the model. In general. releasing code and data is often one good way to accomplish this, but reproducibility can also be provided via detailed instructions for how to replicate the results, access to a hosted model (e.g., in the case of a large language model), releasing of a model checkpoint, or other means that are appropriate to the research performed.
  - While NeurIPS does not require releasing code, the conference does require all submissions to provide some reasonable avenue for reproducibility, which may depend on the nature of the contribution. For example
  - If the contribution is primarily a new algorithm, the paper should make it clear how to reproduce that algorithm.
  - If the contribution is primarily a new model architecture, the paper should describe the architecture clearly and fully.
  - If the contribution is a new model (e.g., a large language model), then there should either be a way to access this model for reproducing the results or a way to reproduce the model (e.g., with an open-source dataset or instructions for how to construct the dataset).
  - We recognize that reproducibility may be tricky in some cases, in which case authors are welcome to describe the particular way they provide for reproducibility. In the case of closed-source models, it may be that access to the model is limited in some way (e.g., to registered users), but it should be possible for other researchers to have some path to reproducing or verifying the results.
5. **Open access to data and code** Question: Does the paper provide open access to the data and code, with sufficient instructions to faithfully reproduce the main experimental results, as described in supplemental material? Answer: [Yes] Justification: The code to reproduce the results on both allosteric and Runs N’ poses benchmark will be provided upon the acceptance of this paper. We plan to upload all the data as the static archive on Zenodo with persistent DOI. Guidelines:
  - The answer [N/A] means that paper does not include experiments requiring code.
  - Please see the NeurIPS code and data submission guidelines (https://neurips.cc/public/guides/CodeSubmissionPolicy) for more details.
  - While we encourage the release of code and data, we understand that this might not be possible, so [No] is an acceptable answer. Papers cannot be rejected simply for not including code, unless this is central to the contribution (e.g., for a new open-source benchmark).
  - The instructions should contain the exact command and environment needed to run to reproduce the results. See the NeurIPS code and data submission guidelines (https://neurips.cc/public/guides/CodeSubmissionPolicy) for more details.
  - The authors should provide instructions on data access and preparation, including how to access the raw data, preprocessed data, intermediate data, and generated data, etc.
  - The authors should provide scripts to reproduce all experimental results for the new proposed method and baselines. If only a subset of experiments are reproducible, they should state which ones are omitted from the script and why.
  - At submission time, to preserve anonymity, the authors should release anonymized versions (if applicable).
  - Providing as much information as possible in supplemental material (appended to the paper) is recommended, but including URLs to data and code is permitted.
6. **Experimental setting/details** Question: Does the paper specify all the training and test details (e.g., data splits, hyperparameters, how they were chosen, type of optimizer) necessary to understand the results? Answer: [Yes] Justification: Experiment settings are discussed in Experiment section and full parameter settings are provided in Supplementary Materials. Guidelines:
  - The answer [N/A] means that the paper does not include experiments.
  - The experimental setting should be presented in the core of the paper to a level of detail that is necessary to appreciate the results and make sense of them.
  - The full details can be provided either with the code, in appendix, or as supplemental material.
7. **Experiment statistical significance** Question: Does the paper report error bars suitably and correctly defined or other appropriate information about the statistical significance of the experiments? Answer: [Yes] Justification: All the the standard errors reported in all the tables and figures are estimated from 1,000 bootstrap by default Guidelines:
  - The answer [N/A] means that the paper does not include experiments.
  - The authors should answer [Yes] if the results are accompanied by error bars, confidence intervals, or statistical significance tests, at least for the experiments that support the main claims of the paper.
  - The factors of variability that the error bars are capturing should be clearly stated (for example, train/test split, initialization, random drawing of some parameter, or overall run with given experimental conditions).
  - The method for calculating the error bars should be explained (closed form formula, call to a library function, bootstrap, etc.)
  - The assumptions made should be given (e.g., Normally distributed errors).
  - It should be clear whether the error bar is the standard deviation or the standard error of the mean.
  - It is OK to report 1-sigma error bars, but one should state it. The authors should preferably report a 2-sigma error bar than state that they have a 96% CI, if the hypothesis of Normality of errors is not verified.
  - For asymmetric distributions, the authors should be careful not to show in tables or figures symmetric error bars that would yield results that are out of range (e.g., negative error rates).
  - If error bars are reported in tables or plots, the authors should explain in the text how they were calculated and reference the corresponding figures or tables in the text.
8. **Experiments compute resources** Question: For each experiment, does the paper provide sufficient information on the computer resources (type of compute workers, memory, time of execution) needed to reproduce the experiments? Answer: [Yes] Justification: Compute overhead and runtimes are discussed in Supplementary Materials. Guidelines:
  - The answer [N/A] means that the paper does not include experiments.
  - The paper should indicate the type of compute workers CPU or GPU, internal cluster, or cloud provider, including relevant memory and storage.
  - The paper should provide the amount of compute required for each of the individual experimental runs as well as estimate the total compute.
  - The paper should disclose whether the full research project required more compute than the experiments reported in the paper (e.g., preliminary or failed experiments that didn’t make it into the paper).
9. **Code of ethics** Question: Does the research conducted in the paper conform, in every respect, with the NeurIPS Code of Ethics https://neurips.cc/public/EthicsGuidelines? Answer: [Yes] Justification: We have reviewed the Code of Ethics and this work conforms. Guidelines:
  - The answer [N/A] means that the authors have not reviewed the NeurIPS Code of Ethics.
  - If the authors answer [No], they should explain the special circumstances that require a deviation from the Code of Ethics.
  - The authors should make sure to preserve anonymity (e.g., if there is a special consideration due to laws or regulations in their jurisdiction).
10. **Broader impacts** Question: Does the paper discuss both potential positive societal impacts and negative societal impacts of the work performed? Answer: [Yes] Justification: See Impact Statement. Guidelines:
  - The answer [N/A] means that there is no societal impact of the work performed.
  - If the authors answer [N/A] or [No], they should explain why their work has no societal impact or why the paper does not address societal impact.
  - Examples of negative societal impacts include potential malicious or unintended uses (e.g., disinformation, generating fake profiles, surveillance), fairness considerations (e.g., deployment of technologies that could make decisions that unfairly impact specific groups), privacy considerations, and security considerations.
  - The conference expects that many papers will be foundational research and not tied to particular applications, let alone deployments. However, if there is a direct path to any negative applications, the authors should point it out. For example, it is legitimate to point out that an improvement in the quality of generative models could be used to generate Deepfakes for disinformation. On the other hand, it is not needed to point out that a generic algorithm for optimizing neural networks could enable people to train models that generate Deepfakes faster.
  - The authors should consider possible harms that could arise when the technology is being used as intended and functioning correctly, harms that could arise when the technology is being used as intended but gives incorrect results, and harms following from (intentional or unintentional) misuse of the technology.
  - If there are negative societal impacts, the authors could also discuss possible mitigation strategies (e.g., gated release of models, providing defenses in addition to attacks, mechanisms for monitoring misuse, mechanisms to monitor how a system learns from feedback over time, improving the efficiency and accessibility of ML).
11. **Safeguards** Question: Does the paper describe safeguards that have been put in place for responsible release of data or models that have a high risk for misuse (e.g., pre-trained language models, image generators, or scraped datasets)? Answer: [N/A] Justification: No risks. Guidelines:
  - The answer [N/A] means that the paper poses no such risks.
  - Released models that have a high risk for misuse or dual-use should be released with necessary safeguards to allow for controlled use of the model, for example by requiring that users adhere to usage guidelines or restrictions to access the model or implementing safety filters.
  - Datasets that have been scraped from the Internet could pose safety risks. The authors should describe how they avoided releasing unsafe images.
  - We recognize that providing effective safeguards is challenging, and many papers do not require this, but we encourage authors to take this into account and make a best faith effort.
12. **Licenses for existing assets** Question: Are the creators or original owners of assets (e.g., code, data, models), used in the paper, properly credited and are the license and terms of use explicitly mentioned and properly respected? Answer: [Yes] Justification: All the datasets and related codes are cited in the main text and Supplementary Materials for further details. Guidelines:
  - The answer [N/A] means that the paper does not use existing assets.
  - The authors should cite the original paper that produced the code package or dataset.
  - The authors should state which version of the asset is used and, if possible, include a URL.
  - The name of the license (e.g., CC-BY 4.0) should be included for each asset.
  - For scraped data from a particular source (e.g., website), the copyright and terms of service of that source should be provided.
  - If assets are released, the license, copyright information, and terms of use in the package should be provided. For popular datasets, paperswithcode.com/datasets has curated licenses for some datasets. Their licensing guide can help determine the license of a dataset.
  - For existing datasets that are re-packaged, both the original license and the license of the derived asset (if it has changed) should be provided.
  - If this information is not available online, the authors are encouraged to reach out to the asset’s creators.
13. **New assets** Question: Are new assets introduced in the paper well documented and is the documentation provided alongside the assets? Answer: [N/A] Justification: All the experiments are evaluated on the publicly available datasets. Guidelines:
  - The answer [N/A] means that the paper does not release new assets.
  - Researchers should communicate the details of the dataset/code/model as part of their submissions via structured templates. This includes details about training, license, limitations, etc.
  - The paper should discuss whether and how consent was obtained from people whose asset is used.
  - At submission time, remember to anonymize your assets (if applicable). You can either create an anonymized URL or include an anonymized zip file.
14. **Crowdsourcing and research with human subjects** Question: For crowdsourcing experiments and research with human subjects, does the paper include the full text of instructions given to participants and screenshots, if applicable, as well as details about compensation (if any)? Answer: [N/A] Justification: This paper doesn’t involve any crowdsourcing or human subjects. Guidelines:
  - The answer [N/A] means that the paper does not involve crowdsourcing nor research with human subjects.
  - Including this information in the supplemental material is fine, but if the main contribution of the paper involves human subjects, then as much detail as possible should be included in the main paper.
  - According to the NeurIPS Code of Ethics, workers involved in data collection, curation, or other labor should be paid at least the minimum wage in the country of the data collector.
15. **Institutional review board (IRB) approvals or equivalent for research with human subjects** Question: Does the paper describe potential risks incurred by study participants, whether such risks were disclosed to the subjects, and whether Institutional Review Board (IRB) approvals (or an equivalent approval/review based on the requirements of your country or institution) were obtained? Answer: [N/A] Justification: This paper doesn’t involve any crowdsourcing or human subjects. Guidelines:
  - The answer [N/A] means that the paper does not involve crowdsourcing nor research with human subjects.
  - Depending on the country in which research is conducted, IRB approval (or equivalent) may be required for any human subjects research. If you obtained IRB approval, you should clearly state this in the paper.
  - We recognize that the procedures for this may vary significantly between institutions and locations, and we expect authors to adhere to the NeurIPS Code of Ethics and the guidelines for their institution.
  - For initial submissions, do not include any information that would break anonymity (if applicable), such as the institution conducting the review.
16. **Declaration of LLM usage** Question: Does the paper describe the usage of LLMs if it is an important, original, or non-standard component of the core methods in this research? Note that if the LLM is used only for writing, editing, or formatting purposes and does *not* impact the core methodology, scientific rigor, or originality of the research, declaration is not required. Answer: [N/A] Justification: LLM is primarily used for writing, editing, formating and data processing. The core methodology does not involve LLM. Guidelines:
  - The answer [N/A] means that the core method development in this research does not involve LLMs as any important, original, or non-standard components.
  - Please refer to our LLM policy in the NeurIPS handbook for what should or should not be described.

## Footnotes

2 As Protenix-v1 does not include MD simulations in the training set, local ensemble generation is inherently unsupported, and, therefore, it was excluded from pose ranking evaluation.

3 Boltz-steering includes various constraints: chirality, ring planarity, internal geometry, steric clashes, overlapping chains

## References

Abramson, J., Adler, J., Dunger, J., Evans, R., Green, T., Pritzel, A., Ronneberger, O., Willmore, K., Ballard, A. J., Bambrick, J., et al. Accurate structure prediction of biomolecular interactions with AlphaFold 3. Nature, 630(8016):493–500, 2024. doi: 10.1038/s41586-024-07487-w. URL https://doi.org/10.1038/s41586-024-07487-w.

Biasini, M., Schmidt, T., Bienert, S., Mariani, V., Studer, G., Haas, J., Johner, N., Schenk, A. D., Philippsen, A., and Schwede, T. Openstructure: an integrated software framework for computa-tional structural biology. Acta Crystallographica Section D: Biological Crystallography, 69(5): 701–709, May 2013. doi: 10.1107/S0907444913007051.

Biswas, A. D., Sabato, E., Vittorio, S., Aletayeb, P., Pedretti, A., Mazzolari, A., Gratteri, C., Beccari, A. R., Talarico, C., and Vistoli, G. Novel method for prioritizing protein binding sites using pocket analysis and md simulations. Heliyon, 11(10):e43084, 2025. ISSN 2405-8440. doi: 10.1016/j.heliyon.2025.e43084. URL https://www.sciencedirect.com/science/article/pii/S2405844025014653.

Buttenschoen, M., Morris, G. M., and Deane, C. M. Posebusters: Ai-based docking methods fail to generate physically valid poses or generalise to novel sequences. Chemical Science, 15(9): 3130–3139, 2024. ISSN 2041-6539. doi: 10.1039/d3sc04185a. URL http://dx.doi.org/10.1039/D3SC04185A.

Ciulli, A. Biophysical Screening for the Discovery of Small-Molecule Ligands, pp. 357–388. Humana Press, Totowa, NJ, 2013. ISBN 978-1-62703-398-5. doi: 10.1007/978-1-62703-398-5_13. URL https://doi.org/10.1007/978-1-62703-398-5_13.

Durairaj, J., Adeshina, Y., Cao, Z., Zhang, X., Oleinikovas, V., Duignan, T., McClure, Z., Robin, X., Studer, G., Kovtun, D., Rossi, E., Zhou, G., Veccham, S., Isert, C., Peng, Y., Sundareson, P., Akdel, M., Corso, G., Stärk, H., Tauriello, G., Carpenter, Z., Bronstein, M., Kucukbenli, E., Schwede, T., and Naef, L. Plinder: The protein-ligand interactions dataset and evaluation resource. bioRxiv, 2024. doi: 10.1101/2024.07.17.603955. URL https://www.biorxiv.org/content/early/2024/07/19/2024.07.17.603955.1.

Isomorphic Labs Team. Accurate predictions of novel biomolecular interactions with isodde. Technical report, Isomorphic Labs, February 2026. URL https://storage.googleapis.com/isomorphiclabs-website-publicartifacts/isodde_technical_report.pdf.

Jumper, J., Evans, R., Pritzel, A., Green, T., Figurnov, M., Ronneberger, O., Tunyasuvunakool, K., Bates, R., Žídek, A., Potapenko, A., et al. Highly accurate protein structure prediction with alphafold. nature, 596(7873):583–589, 2021.

Krivák, R. and Hoksza, D. P2Rank: machine learning based tool for rapid and accurate prediction of ligand binding sites from protein structure. Journal of Cheminformatics, 10(1):39, 2018. doi: 10.1186/s13321-018-0285-8. URL https://doi.org/10.1186/s13321-018-0285-8.

Le Guilloux, V., Schmidtke, P., and Tuffery, P. Fpocket: An open source platform for ligand pocket detection. BMC Bioinformatics, 10:168, 2009. doi: 10.1186/1471-2105-10-168. URL https://doi.org/10.1186/1471-2105-10-168.

Lee, M., Kalicki, C., Jeon, M., Qabel, A., Fadini, A., and AlQuraishi, M. Confornets: Latents-based conformational control in openfold3, 2026. URL https://arxiv.org/abs/2604.18559.

Ma, W., Liu, Z., Yang, J., Lu, C., Zhang, H., and Xiao, W. From dataset curation to unified evaluation: Revisiting structure prediction benchmarks with pxmeter. bioRxiv, 2025. doi: 10.1101/2025.07.17.664878. URL https://www.biorxiv.org/content/early/2025/07/22/2025.07.17.664878.

Masters, M. R., Mahmoud, A. H., and Lill, M. A. Investigating whether deep learning models for cofolding learn the physics of protein-ligand interactions. Nature Communications, 16(1):8854, 2025. doi: 10.1038/s41467-025-63947-5. URL https://doi.org/10.1038/s41467-025-63947-5.

Meller, A., Ward, M. D., Borowsky, J. H., Lotthammer, J. M., Kshirsagar, M., Oviedo, F., Ferres, J. L., and Bowman, G. Predicting the locations of cryptic pockets from single protein structures using the pocketminer graph neural network. Biophysical journal, 122(3):445a, 2023.

Nittinger, E., Özge Yoluk, Tibo, A., Olanders, G., and Tyrchan, C. Co-folding, the future of docking – prediction of allosteric and orthosteric ligands. Artificial Intelligence in the Life Sciences, 8:100136, 2025. ISSN 2667-3185. doi: 10.1016/j.ailsci.2025.100136. URL https://www.sciencedirect.com/science/article/pii/S2667318525000121.

Olanders, G., Testa, G., Tibo, A., Nittinger, E., and Tyrchan, C. Challenge for deep learning: Protein structure prediction of ligand-induced conformational changes at allosteric and orthosteric sites. Journal of Chemical Information and Modeling, 2024.

OpenBind Consortium. Openbind’s first release: A structure–affinity dataset for structure-based ai. https://openbind.uk/news/blog-openbinds-first-release-a-structure-affinity-dataset-for-structure-based-ai/, May 2026. Accessed: 2026-05-07.

OpenFold3 Team. Openfold3-preview2 technical report. Technical report, Open Molecular Software Foundation, 2024. URL https://portal.openfold.omsf.io/reports/of3p2_technical_report.pdf. Accessed: May 2026.

Parikh, V., Foley, B., Gatlin, W., Ludwick, M., Turano, L., and Verkhivker, G. M. Decoding the allosteric paradox: A dual framework integrating ai cofolding models with landscape-guided interpretable ai framework of ligand-protein binding. bioRxiv, 2026. doi: 10.64898/2026.02.24.707829. URL https://www.biorxiv.org/content/early/2026/02/26/2026.02.24.707829.

Passaro, S., Corso, G., Wohlwend, J., Reveiz, M., Thaler, S., Somnath, V. R., Getz, N., Portnoi, T., Roy, J., Stark, H., Kwabi-Addo, D., Beaini, D., Jaakkola, T., and Barzilay, R. Boltz-2: Towards accurate and efficient binding affinity prediction. bioRxiv, 2025. doi: 10.1101/2025.06.14.659707. URL https://www.biorxiv.org/content/early/2025/06/18/2025.06.14.659707.

Prat, A., Zhang, L., Deane, C. M., Teh, Y. W., and Morris, G. M. Sigmadock: Untwisting molecular docking with fragment-based se(3) diffusion, 2026. URL https://arxiv.org/abs/2511.04854.

Purnomo, J., Sun, K., and Head-Gordon, T. Cafe: A co-folding approach for fragment exploration of allosteric and cryptic binding sites. bioRxiv, pp. 2026–07, 2026.

Richman, D. D., Karaguesian, J., Suomivuori, C.-M., and Dror, R. O. Unlocking hidden biomolecular conformational landscapes in diffusion models at inference time, 2026. URL https://arxiv.org/abs/2512.03312.

Shi, Y. A glimpse of structural biology through x-ray crystallography. Cell, 159(5):995–1014, 2014. ISSN 0092-8674. doi: 10.1016/j.cell.2014.10.051. URL https://www.sciencedirect.com/science/article/pii/S0092867414014238.

Singhal, R., Horvitz, Z., Teehan, R., Ren, M., Yu, Z., McKeown, K., and Ranganath, R. A general framework for inference-time scaling and steering of diffusion models, 2025. URL https://arxiv.org/abs/2501.06848.

Škrinjar, P., Eberhardt, J., Tauriello, G., Schwede, T., and Durairaj, J. Have protein-ligand cofolding methods moved beyond memorisation? bioRxiv, 2025a. doi: 10.1101/2025.02.03.636309. URL https://www.biorxiv.org/content/early/2025/08/04/2025.02.03.636309.

Škrinjar, P., Eberhardt, J., Tauriello, G., Schwede, T., and Durairaj, J. Have protein-ligand cofolding methods moved beyond memorisation? bioRxiv, 2025b. doi: 10.1101/2025.02.03.636309. URL https://www.biorxiv.org/content/early/2025/08/04/2025.02.03.636309.

Stepniewska-Dziubinska, M. M., Zielenkiewicz, P., and Siedlecki, P. Improving detection of proteinligand binding sites with 3d segmentation. Scientific Reports, 10(1):5035, 2020. doi: 10.1038/s41598-020-61860-z. URL https://doi.org/10.1038/s41598-020-61860-z.

team, C. D., Boitreaud, J., Dent, J., McPartlon, M., Meier, J., Reis, V., Rogozhonikov, A., and Wu, K. Chai-1: Decoding the molecular interactions of life. bioRxiv, 2024. doi: 10.1101/2024.10.10.615955. URL https://www.biorxiv.org/content/early/2024/10/11/2024.10.10.615955.

Team, G. R., Dobles, A., Jovic, N., Leidal, K., Murugan, P., Williams, D. C., Wulsin, D., Gruver, N., Ji, C. X., Pruegsanusak, K., Scarpellini, G., Sharma, A., Swiderski, W., Bootsma, A., Bowen, R. S., Chen, C., Chen, J., Dämgen, M. A., DiFrancesco, B., Fishman, J. D., Ivanova, A., Kagin, Z., Li-Bland, D., Liu, Z., Morozov, I., Ouyang-Zhang, J., Iv, F. C. P., Shah, K. S., Shor, B., da Silva, G. M., Tal, R., Tessmer, M., Tilbury, C., Vetcher, C., Zeng, D., Al-Shedivat, M., Faust, A., Feinberg, E. N., LeVine, M. V., and Pan, M. Pearl: A foundation model for placing every atom in the right location, 2025. URL https://arxiv.org/abs/2510.24670.

Team, P., Zhang, Y., Gong, C., Zhang, H., Ma, W., Liu, Z., Chen, X., Guan, J., Wang, L., Yang, Y., Xia, Y., and Xiao, W. Protenix-v1: Toward high-accuracy open-source biomolecular structure prediction. bioRxiv, 2026. doi: 10.64898/2026.02.05.703733. URL https://www.biorxiv.org/content/early/2026/02/22/2026.02.05.703733.1.

Trott, O. and Olson, A. J. AutoDock Vina: improving the speed and accuracy of docking with a new scoring function, efficient optimization, and multithreading. Journal of Computational Chemistry, 31(2):455–461, 2010. doi: 10.1002/jcc.21334.

Wang, J. and Dokholyan, N. V. Unified protein-small molecule graph neural networks for binding site prediction. bioRxiv, 2025. doi: 10.1101/2025.09.03.674017. URL https://www.biorxiv.org/content/early/2025/09/08/2025.09.03.674017.

Xie, J., Wang, S., Xu, Y., Deng, M., and Lai, L. Uncovering the dominant motion modes of allosteric regulation improves allosteric site prediction. Journal of Chemical Information and Modeling, 62 (1):187–195, 2022. doi: 10.1021/acs.jcim.1c01267.

Yu, H., Bekar-Cesaretli, A. A., Lazou, M., Kozakov, D., Joseph-McCarthy, D., and Vajda, S. Bias in the AlphaFold3 prediction of ligand-induced domain motion in enzymes. Proceedings of the National Academy of Sciences, 123(10):e2530709123, 2026. doi: 10.1073/pnas.2530709123. URL https://doi.org/10.1073/pnas.2530709123.

Zhang, Y., Gong, C., Sun, J., Guan, J., Ren, M., Xue, S., Zhang, H., Ma, W., Liu, Z., Chen, X., and Xiao, W. Protenix-v2: Broadening the reach of structure prediction and biomolecular design. bioRxiv, 2026. doi: 10.64898/2026.04.10.717613. URL https://www.biorxiv.org/content/early/2026/04/11/2026.04.10.717613.

Zhao, L., Zhu, Y., Wang, J., Wen, N., Wang, C., and Cheng, L. A brief review of protein–ligand interaction prediction. Computational and Structural Biotechnology Journal, 20:2831–2838, 2022. doi: 10.1016/j.csbj.2022.06.004.

